# Persistence of neuronal representations through time and damage in the hippocampus

**DOI:** 10.1101/559104

**Authors:** Walter G. Gonzalez, Hanwen Zhang, Anna Harutyunyan, Carlos Lois

## Abstract

It is not known how neurons encode memories that can persist up to decades. To investigate this question, we performed simultaneous bilateral imaging of neuronal activity in the mouse hippocampus over weeks. From one day to the next ∼40 % of neurons changed their responsiveness to cues; however, thereafter only 1 % of cells changed for each additional day. Despite the apparent instability between days, field responses of CA1 neurons are very resilient to lack of exposure to the task or lesions to CA1. Although a small fraction of individual neurons retain their responses over weeks, groups of neurons with inter- and intrahemispheric synchronous activity had stable responses. Neurons whose activity was synchronous with a large group of neurons were more likely to preserve their responses across multiple days. These results suggest that although information stored in individual neurons is relatively labile, it is very stable in networks of synchronously active neurons.

**One Sentence Summary:** Neuronal representations in networks of neurons with synchronized activity are stable over weeks, even after lack of training or following damage.

## Main Text

Memories are processed and stored by a complex network of neurons across several circuits in the brain; however, little is known about how information is encoded and retained in these neurons for long periods of time. Information could be stored at different hierarchical levels, within individual neurons through modification of their synapses, or distributed among many neurons in different brain areas including the hippocampus and cortex. The hippocampus is known to play an essential role in the formation of memories (*1, 2*) and neurons in this brain area show robust response to space (place cells), time (time cells), or other task relevant cues (*3–5*). Many works have studied how neuronal activity in the hippocampus changes during learning (*6, 7*), attention (*8, 9*), and re-exposure (*10*). However, what aspects of neuronal activity in the hippocampus persist during future visits to a familiar environment, how is information encoded in group of neurons, and how lesions perturb the long-term maintenance of these neuronal patterns remains poorly understood.

The question of how information is stably encoded in neurons in the hippocampus remains a controversial issue. Whereas extracellular recordings show that place cells retain their fields from days to a weeks (*11*), calcium imaging experiments show drastic changes of neuronal activity across days (*12–15*). Considering that the hippocampus is necessary for the formation, but not for the long-term maintenance of memories, it is possible that neuronal representations in the hippocampus may change over time as information is recalled from other brain areas (*16*). Previous long-term imaging of neuronal activity showed that a large number of pyramidal neurons in CA1 are active during a 35 day period but only 31 % were active on any given session and only 2.8 ± 0.3% were active in all sessions (*12*). However, the inability to detect active neurons on consecutive days could be due to motion artifacts caused by the removal and reattachment of the microendoscope between days. In addition, overexpression of GCaMP using AAVs can induce cell toxicity or even death (*17*). To overcome these potential limitations we built custom microendoscopes that were chronically implanted and performed long-term simultaneous bilateral imaging of hippocampal activity in freely moving Thy1-GCaMP6s mice **(Figure 1a, S1, Movie 1 and 2, see supplementary data)** (*18–21*). The combination of chronic implants, high sensitivity microendoscope, improved cell detection and registration with the CNMFe software allowed us to minimize motion artifacts and to increase the reliability of long-term recordings **(Fig. 1b-c and S2)**(*22, 23*). We imaged CA1 pyramidal neuron activity for several weeks through three situations, which we defined as follows: (i) “learning”, during the initial 5 sessions of exposure to a novel linear track with sugar water reward at the ends, (ii) “re-exposure”, after a 10-day period during which the animal was not exposed to the linear track, and (iii) “damage”, following light-induced hippocampus lesion **(Fig. S1)**. We observed robust single neuron activity across days for up to 8 months **(Fig. 1d-e and Movie 3 and 4).** We did not notice significant differences between hemispheres and unless stated otherwise, the values reported represent combined data from both hemispheres. In any given day, 88 ± 4 % of all neurons in the field of view were active and 51 ± 17 % were active every session (**Fig. 1f-h and S3**). Within a day, the vast majority of neurons (95 %) were active both while mice explored their home cage or while running in the linear track. However, in the same day between environments (cage or linear track) or across days in the linear track, neurons displayed significant changes in their firing rates (**Fig. S4**). Thus, minimizing motion artifacts using chronic implants and improved signal extraction and registration allowed us to observe that most CA1 neurons that are active one day are also active on subsequent days.

**Figure 1.**
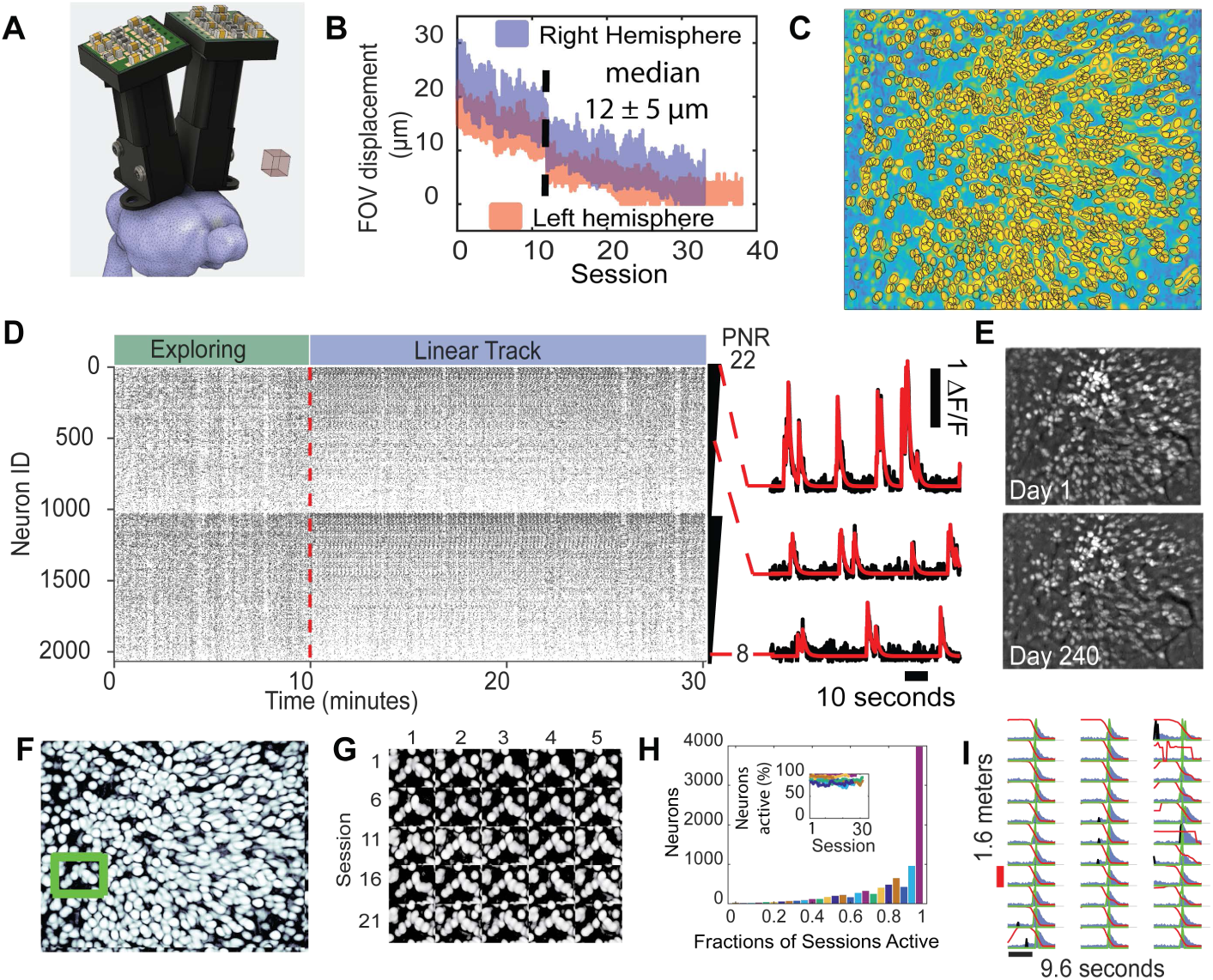
Long-term simultaneous bilateral imaging of CA1 activity in freely moving transgenic mice. **(A)** Rendering of custom endoscope used in the bilateral configuration. **(B)** Chronic implants show small field of view (FOV) displacement across weeks. Dashed line represents period of no task (10 days). **(C)** Multiple sessions (25) combined and analyzed simultaneously using CNMFe (see methods). **(D)** Raster plot of deconvoluted neuronal activity in the right (top) and left (bottom) hemisphere of a trained freely moving mouse. Neurons are ranked from high to low peak-to-noise ratio (PNR). Between 754-1480 neurons were recorded per hemisphere, median 1192, in 4 bilateral mice and 5 with unilateral microendoscopes). Total 15084 neurons, 4401 place/time cells. (**E**) Maximum intensity projection in the field of view in one animal recorded for 8 months. (**F-G**) Intensity correlation map of a single session (*left*) and a small region of interest (green rectangle) showing the persistence of activity throughout 25 sessions (*right*). (**H**) Activity distribution of the right hemisphere showing that the majority of neurons are active on most sessions (n=8 mice). **(I)** Activity of one place cell when the mouse passed its receptive field near the middle of the maze. Blue area represents the deconvoluted calcium transient, green discrete peak is the deconvoluted activity used for calculation of the place field, black deconvoluted activity not included because the animal was not moving. Red trace represents the location of the mouse in the linear track. The first twenty-one consecutive passes shown.

To acquire a comprehensive view of stability of neuronal representations in CA1, we studied both cells that were active in specific locations of the maze when animals were running (defined as “place cells”), and during periods of immobility (defined as “time cells”) (*24, 25*) **(Fig. S5-7)**. Previous work reported that place fields underwent drastic changes across sessions, reaching near random levels (75-85 % change) on the following session (*12*). Under our conditions, we observe that from one day to the next 62 ± 12 % of neurons retained response to a field (defined here as “cell overlap”). These changes were present regardless of how familiar the animal was to the environment but were more prominent during the “learning” phase **(Fig. S5e)**. Surprisingly, despite the initial abrupt change, the cell overlap for subsequent days decreased only an additional ∼1 % per day, reaching random levels after ∼ 50 days (**Fig. 2a-c**).

**Figure 2.**
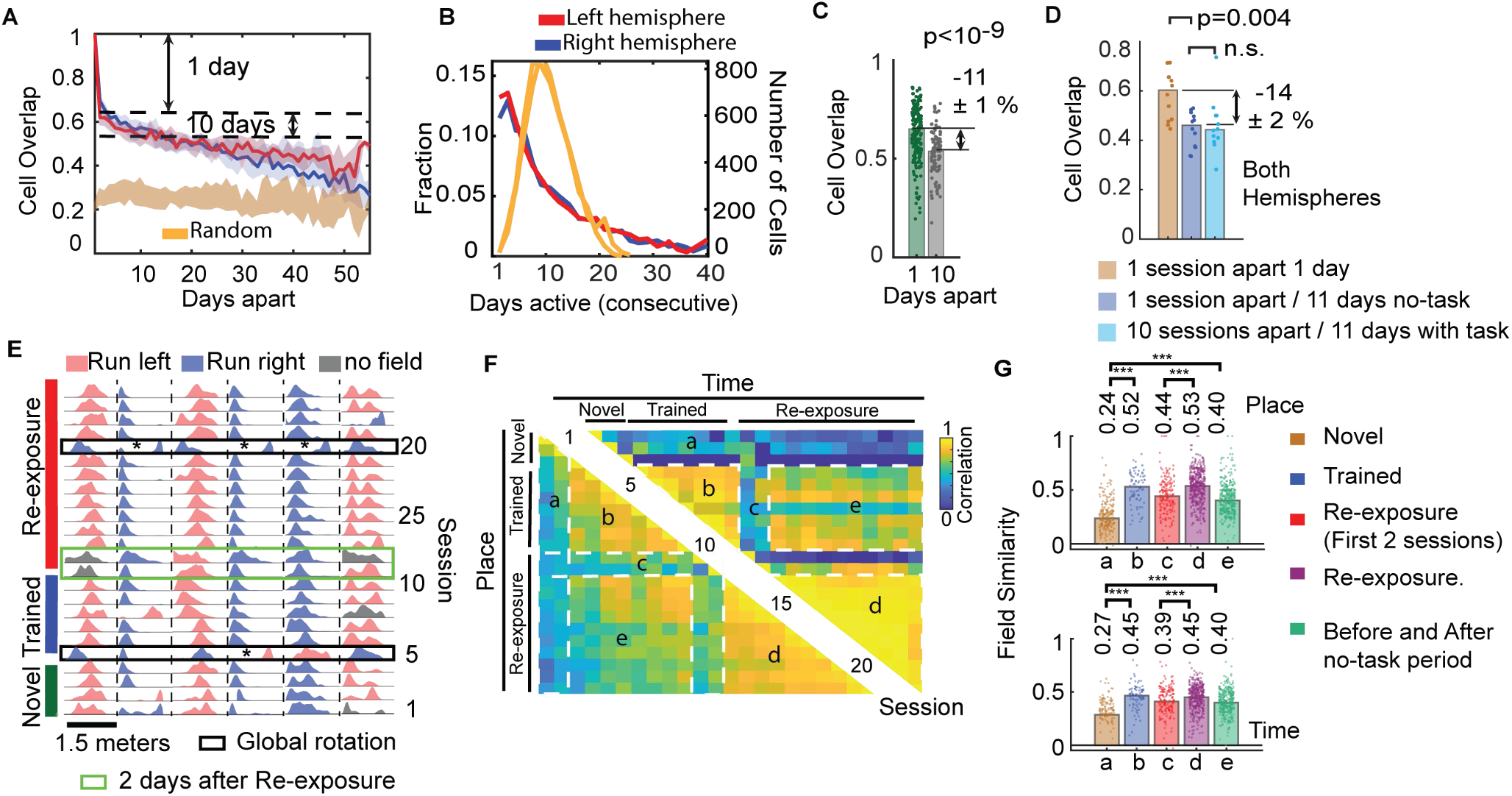
Spatial and temporal representations persist through periods of no exposure to the task. **(A)** The probability of a place/time cell in one session having a field response N-days apart. Trained, re-exposed, and learning periods were included in this analysis. Solid lines represent the median and shadow the 95 % bootstrap confidence interval. Random level represent overlap if neurons randomly became responsive to a field on the following day. **(B)** Distribution of neurons retaining their field response for N consecutive sessions (5 left hemisphere, 8 right hemisphere, median ± std, 10 ± 5 days). **(C)** Quantification of cell overlap 1 day (62 ± 12 %, n = 348 session pairs) and 10 days apart (51 ± 16 %, n = 106 session pairs, ranksum-test). The analysis in (a-c) is between all possible session intervals including the no task period. **(D)** Quantification of cell overlap 1 day apart, 11 days apart without task, and 11 days apart with task (60 ± 12 %, 46 ± 7 %, 44 ± 11 %, respectively, n=12). **(E)** Response fields of six place cells across 35 days. Black rectangles show sessions in which 87 ± 8 % of fields changed direction. Bidirectional cells not marked. Asterisks show place cells with fields whose field position with respect to the reward port remains similar. Green rectangle are the two sessions after re-exposure. **(F)** Pair-wise field correlation of neurons in the right hemisphere. Colors represent the median value. Numbers indicate the session. Sessions were grouped as indicated by the dashed rectangle and letters. Only neurons with field responses on both sessions compared were considered, sessions with rotated representation (20 and 5) were removed for clarity. **(G)** The similarity between fields in each group of sessions was obtained by subtracting the correlation of randomly aligned neurons (n = 7, right hemisphere, each dot is the median between two sessions, p< 10^−4^ marked by ***, sessions with rotated representations not excluded from the analysis). During the first 2 days of re-exposure mouse performance in the track did not degrade (figure S1b).

Repetitive exposure to the task could induce changes in neuronal representation through continuous updates in place and time fields due to minor changes in environment (i.e. different personnel or odors in the room). To investigate this possibility, we introduced a no-task period in which trained animals were not exposed to the linear track for 10 days **(Fig. S1b)**. We then compared the changes in place and time cells between animals not exposed to the track and animals which were continuously exposed to the track. Following re-exposure, place and time cells in sessions separated by 11 days that included the 10-day period of no-task changed their fields by a similar fraction as animals continuously exposed to the task in sessions separated by 11 days (**Fig. 2d and S8**). Thus, changes in place and time cells happen independently of whether the animal is exposed to the task.

Next, we analyzed whether the fields to which a neuron is responsive changes between sessions. Response fields were relatively similar across days (correlation of 0.7 ± 0.3, 505 neurons, during the “trained” period between one to 5 days apart) (**Fig. 2f-g**). Interestingly, in some sessions, we observed that 87 ± 8 % of cells changed the direction of their fields by 180 ° in the linear track, reversing the directional representation encoded in the previous session (**Fig. 2e**). Rotations of fields happened simultaneously across hemispheres and were accompanied by minimal changes in animal behavior (**Fig. S9**). Rotations in the representation of the linear track could explain the observed low stability of place fields during navigation in a virtual track (*14*). Field similarities across days were also significantly lower during periods of learning and during the first two days following re-exposure to the task (**Fig. 2f-g)**. Response fields fluctuate from day to day during the learning periods or immediately following re-exposure (regions a and c in figure 2f), but once the animal becomes familiar with the task, place and time cells can recover their fields even following a 10-day period in which the animal is not exposed to the task (region e in figure 2f).

To investigate whether CA1 representations are also resilient to brain damage we performed local lesions induced by increasing the LED power and illumination time of the implanted microendoscopes (5 to 10-fold over the threshold needed to visualize GCaMP) (**Fig. 3a**). High illumination intensity induces a local increase in the tissue temperature and affects neurons along a spectrum ranging from perturbing their firing activity to triggering their death. One day after light damage, we observed a massive increase in synchronized firing analogous to interictal discharges (**Fig. 3b**). These abnormal bursts of activity recruited the majority of neurons in the field of view and were direction specific in the linear track **(Fig. S10b and Movie 5)**. During days with abnormal CA1 activity, the firing behavior of neurons changed dramatically, and the number of place and time cells increases significantly (**Fig. 3c and S10e**). However, during this period, place and time field correlation across days decreased to near random levels **(Fig. 3d-f)**. Interestingly, after 2 to 10 days, abnormal activity ceased overnight but not all neurons retained their activity in the linear track **(Movie 4)**. In a similar fashion to what we observed in the re-exposure experiments, after recovery from damage, place and time fields stabilized and a significant portion of place and time cells were responsive to the same fields that they had before the lesion (81 ± 11 % of cells with a field before and after the lesion had a correlation above 0.7, compared to random p<10^−10^ ranksum, n = 49 sessions) **(Fig. 3f)**.

**Figure 3.**
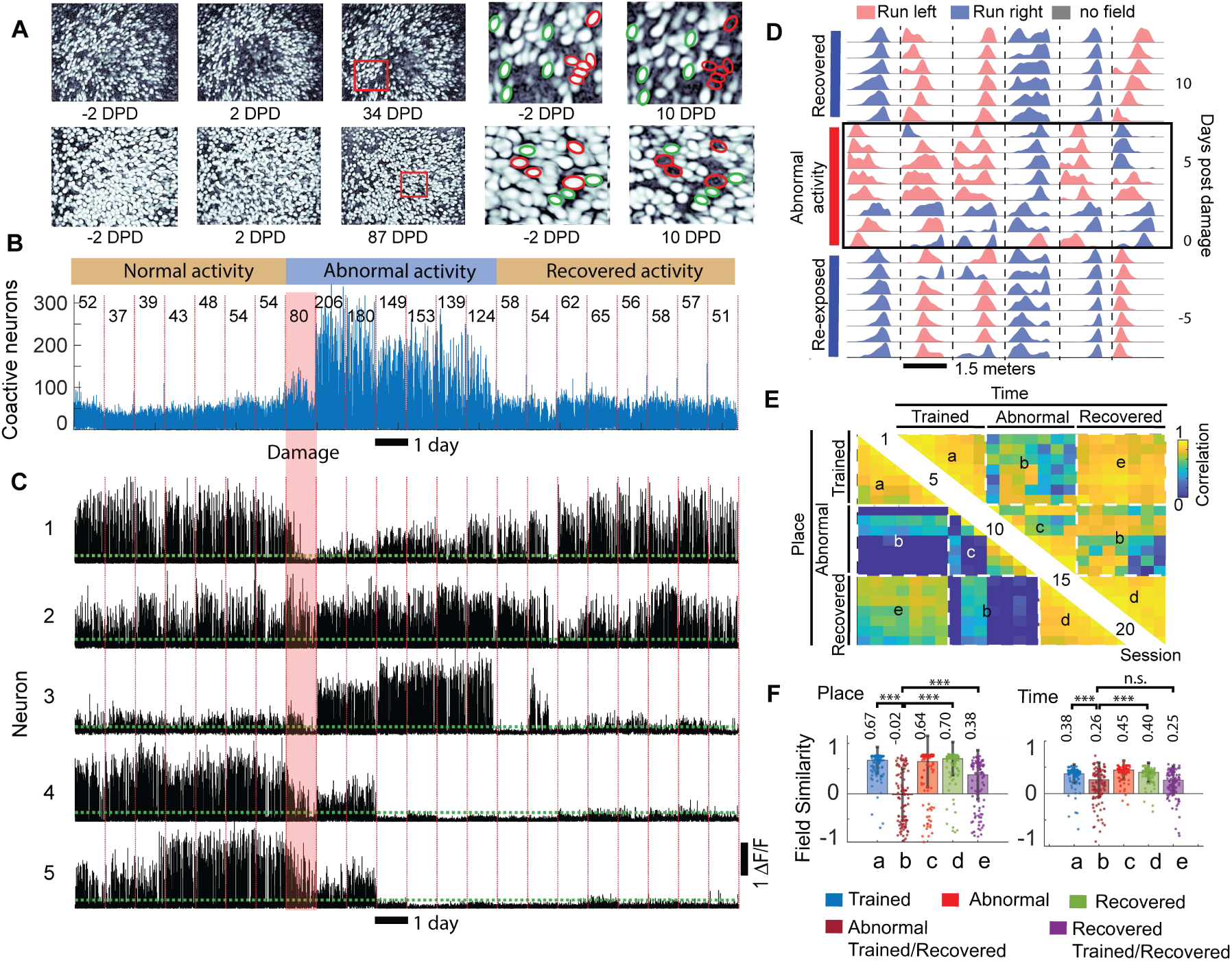
Tissue heating induces direction specific bursting activity and unstable fields followed by partial recovery of the original representation. **(A)** Intensity correlation of a 20-minute session before, and after a lesion induced by high illumination intensity (DPD, days post damage). Top row represents one animal with a large lesion and the right two panels show some neurons that remain active (green circle) or become inactive (red circle) after the lesion. The bottom panel shows another animal with a smaller lesion. Both animals had a clear direction dependent abnormal activity in the linear track (**Movie 5 and 6**). **(B)** Number of neurons active simultaneously (within 160 milliseconds) during each session (number shows median of the largest 90% bursts within a day). Each day is marked by a vertical red line. **(C)** Single neuron activity across days while the mouse runs in the linear track (black traces) showing changes in activity induced by tissue damage (red rectangle). The horizontal green line shows the three-standard deviation of the background fluctuations. Top two neurons recover from the damage, middle neuron is highly active during abnormal activity, and the last two neurons become inactive following damage. **(D)** Activity distribution of six place cells showing the high instability of fields during abnormal activity, colors as in figure 2. Black rectangle shows sessions with abnormal activity. **(E)** Pair-wise correlation between sessions of neurons which retained their fields (right hemisphere). **(F)** The similarity between fields in groups of sessions before, during, and after the lesion (n = 2 mice, each dot is the median of a session pair, numbers are median, p< 10^−4^ marked by ***).

It has been suggested that groups of neurons with synchronized activity form cell assemblies able to encode learned representations for long periods of time. Cell assemblies encoding temporal and spatial cues have been observed in the hippocampus (*26, 27*). However, whether these assemblies develop during learning, how many neurons participate in them, for how many days they persist, and whether they can encode stable information across days is not known. We started analyzing the activity of pairs of neurons, the simplest level of synchronized groups of cells. The Pearson’s correlation between neurons is independent of the firing rate; however, we observe that neurons in CA1 become more synchronized with other neurons within and across hemispheres as mice become familiar with the linear track (**Fig 4a**). The increase in synchronous pairs arises mainly due to the activity of place and time cells and is proportional their firing rate **(Fig. S11-12)**. Synchronized pairs of neurons also tended to make the same errors as the animal performed the task, as illustrated in two scenarios. First, whenever one neuron in the pair failed to fire in its field, 50 ± 30 % of the time the other neuron in the pair failed as well. Second, when pair of neurons fired, their deviation from the field peak was highly correlated (0.71 ± 0.14, p<10^−8^) (**Fig. S12e**). Across days, we observed that the likelihood of a neuron maintaining its responsiveness to a field was proportional to the degree of synchrony it had with another neuron in a pair **(Fig. S12g)**. Altogether, these results support the notion that synchronized activity does not occur simply by chance due to overlapping fields, and it may be responsible for the stability of representations over time.

**Figure 4.**
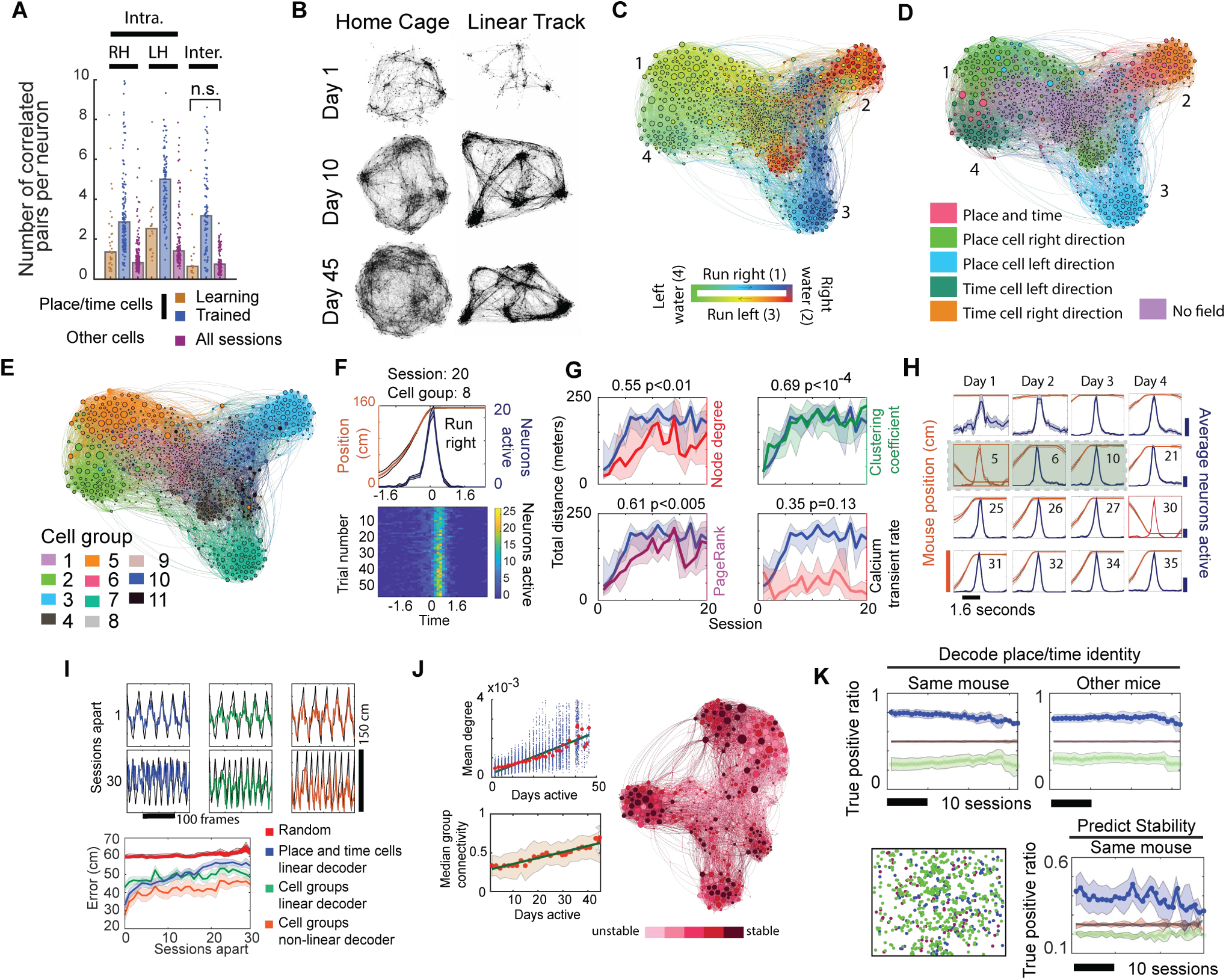
Synchronous activity in group of cells encode stable representations of the task. **(A)** Number of inter- and intra-hemispheric synchronous pairs as a function of training (4 bilateral mice, each dot represents a session in one mouse, all differences are significant p<10^−6^). **(B)** Graph topology in the linear track and home cage of one mouse during learning, trained, and re-exposed periods (data from one hemisphere used, lines represent correlation > 0.10, neurons shown as small dots). **(C)** Graph colored by the median location where each neuron (node) was active during a 20-minute session in the linear track (right hemisphere). **(D)** Graph of neuronal activity in the linear track colored by experimentally determined field responses. **(E)** Network graph colored by cell group extracted by the Markov diffusion approach. The size of each node in (c-e) corresponds to the degree (number of connecting lines) of each node. **(F)** (Top panel) Integrated activity of neurons in a cell group during a linear track session and (bottom panel) activity each time the mouse ran through the cell group’s receptive field. **(G)** Graph topology evolve on timescales of days and are correlated with task performance. The increase in node degree and PageRank during learning indicate that neurons become synchronized and some neurons have a higher importance in the network than others. The development of cell groups during learning induces an increase in the clustering coefficient whereas no relationship between firing rate and behavior is observed. Solid lines represent the median of the correlation between graph metric and behavior, the shadow is the 95 % bootstrap confidence interval (n=12). **(H)** Persistence of a cell group identified by neuronal activity from days 5-6 (shadows represent deviation during a session). Activity of one cell group is shown across 35 days (number indicates day, red traces are sessions with rotated representations). Shaded figures show the sessions used to extract the cell group (see methods). **(I)** Mouse position in the linear track decoded using all place/time cells (132 to 639 neurons), or the summed activity of all neurons in 11 cell groups. The performance of decoders using cell group activity outperformed place/time cell decoders on long-timescales (20+ sessions apart, random vs place/time linear decoder p=0.210 and random vs cell group linear decoder, p<10^−5^, Friedman ANOVA). Solid lines represent the median ± std. of 12 mice. Only periods when the mice were moving are shown but they are included in the calculations of the errors. **(J)** Graph colored by the fraction of sessions a neuron was classified as a time or place cell, larger nodes have larger number of synchronous pairs (degrees). The top inset shows the relationship between degree of a neuron and on how many days that neuron will be responsive to a field (red dots mark median or all nodes, solid line is a linear fit to the blue dots, R-square 0.82, p<10^−10^, one-way ANOVA). The connectivity of a neuron within the cell group it belonged was also proportional to its stability across days (R-squared 0.91, p<10^−17^, one-way ANOVA) and correlated to the Shannon information content of the neuron (0.52 ± 0.26, p<0.01, two-sided T-test). **(K)** (Left) A decoder trained with graph topology metrics on one day can identify 71 ± 11 % of all place/time cells the next day and 59 ± 10 % after 30 days in the same mouse. (Right) Similar results are obtained when a decoder trained with one mouse is used to identify place/time cells in other mice (n = 7). The decoders also identify 86 ± 2 % of neurons with no field responses. (Bottom Right) Neurons that will become responsive or unresponsive to a field between two sessions apart can be decoded from graph metrics 46 ± 5 % of the time. The bottom left figure shows neurons identified as place/time cell (green dots), not identified (blue dots), or misclassified (red dots) in the same mouse.

To explore this hypothesis, we analyzed correlations of neuronal activity to identify whether neuron pairs belong to a larger cell assembly and whether stable information could be encoded in these larger neuronal networks (*26*). Network graphs where links are defined based on correlation, show a clear behavior-dependent topology, evolving throughout periods of learning and undergoing extensive reorganization upon transition from exploring in the home cage to running in the linear track **(Fig. 4b)**. Graphs revealed the presence of dense clusters comprised of neurons within and across hemispheres with preferences for specific behaviors **(Fig. 4c-d, S13 and Movie 7)**. Extraction of these clusters using a Markov diffusion approach identified groups of neurons (defined here as cell groups) encoding direction-specific information about several aspects of the task, including periods of running, immobility, drinking, and turning (**Fig. 4e-f and Movie 8-9**) (*28*). Synchronized activity of cell groups was specific for the environment to which the animal was exposed. Cell groups that had synchronous activity in the linear track became asynchronous in the home cage, and vice versa (**Fig. S13d**). Individual neurons developed their responsiveness to a field within minutes of exposure to the track. In contrast, the functional connectivity of neurons in a graph and neuron clustering develop over 2-4 sessions **(Fig. 4g)**. Indicating that neuronal synchrony does not arise simply by field overlap and highlights a reorganization of network activity during learning which is highly correlated with task performance (clustering coefficient correlation of 0.69, p<10^−4^, n=12). Moreover, the task information encoded in these groups did not degrade over time (up to 35 days), even after a 10-day period of no task (**Fig. 4g-h**). Thus, using features in the correlation matrix of neuronal activity we demonstrate that groups of neurons in CA1 can encode persistent representations of the task across weeks even if the activity of individual neurons varies over time.

The analysis of correlated neuronal activity of CA1 neurons has been used to decode the behavior of the animal or the response of neurons to a field (*29, 30*), however, it is not known whether this activity can be used to predict whether a cell will be responsive to a field into the future (across days). Using graphs, we observed that the likelihood that a neuron would maintain its responsiveness to a field over multiple days was proportional to the extent of connectivity within the cell group it belonged to (**Fig. 4i**). The Shannon information content of neurons was also proportional to how synchronized a neuron was with other cells with similar response fields (correlation 0.52 ± 0.26, p<0.01). To test the hypothesis that neuronal correlation can be used to predict the responsiveness of a neuron to a field we analyzed metrics describing graph and node connectivity and trained a decoder to determine whether a neuron would be a place or time cell in the future (see methods). Using this approach, we show that the synchrony-stability relationship can be used to decode which neurons in a session will be responsive to a field and predict whether a neuron will maintain its field N sessions apart, even after a period of 10 to 20 days **(Fig. 4k, see methods)**. Furthermore, a decoder trained to identify place and time cells from graph metrics in one trained animal in one session can identify place and time cells in other animals simply by analyzing the correlation matrix of neuronal activity. Thus, the features of a neuron in a graph are sufficient to decode signatures that are specific to time cells, specific to place cells, or even specific for cells that are neither time or place cells, and these signatures are common between animals.

In contrast to previous studies on the stability of CA1 representations, we observed that the vast majority of neurons are active on most days but their firing rate changes across sessions and tasks (*12*). We observe high stability when analyzing the cell overlap of place and time cells such that even after 35 days 40 % of neurons were responsive to a field (*12, 14*). Our results indicate that hippocampal representations change drastically from one day to the next, but much more slowly thereafter (an additional ∼1% change per day). In addition, we found that the representations in CA1 were able to recover after an extended period (10 days) without performing the task, or even after abnormal activity induced by local lesions. These manipulations revealed a common feature of information persistence. In both cases, fields undergo transient drifts and fluctuation ultimately converging to a neuronal representation similar to that present before perturbations. These findings provides strong experimental evidence for the presence of attractor-like ensemble dynamics as a mechanism by which the representations of an environment are encoded in the hippocampus (*31*). Interestingly, spatially triggered hippocampal discharges were observed in transgenic mice with chronic blockade of CA2 synaptic transmission (*32*). Our results show that such phenotype can be elicited by unilateral lesions to CA1 and more importantly, CA1 can recover from such abnormal activity. These results suggest a model with two complementary features. First, neuronal representations spontaneously change over time, such that cells whose fields persist longer than 35 days are rare. Second, there are mechanisms that ensure the persistence of representations over short periods of time (days) even if the animals are not training in the task, or if the circuit is perturbed by lesions.

The results presented here reveal a more heterogeneous organization of information in CA1 during learning, recall, and following recovery from damage. When naïve mice are first exposed an unfamiliar environment, place and time cells become sensitive to a field, but their response is stochastic and unstable between days. As mice become familiar with the task and environment, place/time cells become synchronized within and between hemispheres, form cell groups, become more stable between days and resilient to perturbations. At the ensemble level, cell groups form stable representation of the environment but neurons in these cell groups are not all equally stable. Between any given day, ∼ 40 % of cells change their field response; however, on average individual neurons can retain their field response for 10 ± 5 consecutive days, with an even smaller number being active for over 4 weeks. These results are analogous to studies in zebra finches showing that highly stereotypic behaviors are characterized by unstable participation of excitatory neurons but stable ensemble activity (*33*). Analysis of neuronal correlation using graphs reveals that these heterogeneities are related to how synchronous a neuron is with other neurons encoding similar information, such that neurons more connected to their cell group become more stable and have higher information content than neurons at the boundary of a cell group. Lastly, we show that the features of a neuron in a graph are also sufficient to decode whether neurons encode information about the task without knowledge of the animal’s behavior. Overall, our findings suggest a model where the patterns of activity of individual neurons gradually change over time while the activity of groups of synchronously active neurons ensures the persistence of representations.

## Supplementary Material

### Methods

#### Animals

Male and female C57BL6J-Tg-Thy1-GCaMP6s 6 to 20-week old (Jackson Labs stock: 025776) were housed in a reverse 12 h light/dark photocycle and provided food and water ad libitum. Mice were single housed post-surgery until the end of the experiment. Experimental animals were selected randomly and include both sexes. All animal procedures were approved and performed following institutional guidelines (Caltech IACUC).

#### Unilateral/Bilateral endoscope implantation

Mice were anesthetized with a single dose of 100/10 mg/kg ketamine/xylazine before the surgery and placed into a stereotactic frame. The body temperature was maintained with a passive heating pad at 37 °C. Ketoprofen 5 mg/kg and buprenorphine SR 1 mg/kg was subcutaneously injected prior to surgery. Bupivacaine 1 mg/kg solution added dropwise along the surgical incision prior to wound closure and animals were maintained on ibuprofen 30 mg/mL (in the water) ad libitum for at least 3 days post-surgery. Animals were in a recovery period for at least 4 weeks before attachment of the microendoscope.

Mice underwent unilateral or bilateral surgeries to implant 1.8 mm GRIN lenses directly dorsal to CA1. Before implantation, we performed a 1.8 mm diameter craniotomy centered around the coordinates (relative to bregma: 1.8 mm and −1.8 mm lateral; −2.0 mm posterior) using a FG1/4 carbide bur. Freshly prepared artificial cerebrospinal fluid (aCSF) was applied to the exposed tissue throughout the surgery to prevent dehydration. Using a blunt 26-gauge needle, the dura, cortex, and portion of the corpus callosum were quickly aspirated under continuous perfusion with aCSF. Aspiration was stopped once a thin layer of horizontal fibers was left on the surface of the hippocampus. The cortical cavity was perfused with more aCSF and small pieces of moist gelfoam were placed on the surface of the craniotomy to prevent excessive bleeding while avoiding contact with the surface of hippocampus. Once the surface of the hippocampus was clear of blood, the GRIN lens was slowly lowered in into the brain using a stereotaxic arm to a depth of 1.30 mm below the surface of the skull. Removal of the cortex and insertion of the GRIN lens was performed in 10 minutes or less (in each hemisphere) to prevent bulging of the hippocampus due to the decreased dorsal pressure. Two skull screws were placed anterior to bregma (1.8/-1.8 mm lateral; 1.0 mm anterior) and both the screws and lens were secured with cyanoacrylate and dental cement. The exposed end of the GRIN lens was protected with transparent Kwik-seal glue and animals were returned to a clean cage. Two weeks after the surgery, mice were anesthetized with 1.0 to 2.0 % isoflurane, the glue covering the GRIN lens was removed and a microendoscope was aligned with the GRIN lens. The miniature microscopes were connected to a portable computer which provided live view of the fluorescence image though the endoscopic lens and guided the final alignment and focal plane of the microscope and lens. The microscope was permanently attached to the implant with dental acrylic and the focal sliding mechanism on the microendoscope was sealed with superglue.

#### Local CA1 damage

Damage was induced unilaterally by increasing the LED power of the microendoscope to the maximum allowable level. The power of the 470 nm blue LED on the miniscope was measured to produce 500-700 μW at this setting, measured at the end of the GRIN lens facing CA1, using a power meter (Thorlabs PM100D, S155C probe). The brain was illuminated for at least 30 minutes during the foraging and linear track task. In one animal we performed this heating session on two sessions in order to increase the damage area (top row of figure 3a). The extent of the lesion was assessed by the presence of abnormal burst of activity and by the presence of continuously green cells on the following day. Sessions with abnormal activity were selected based on the low place/time field correlation observed in figure 3e and by the increase number of co-active neurons, as shown in figure 3b.

#### Mouse behavior

Mice were maintained in a reverse photo cycle and three days prior to initialization of behavior recording they were set under a water restriction protocol in which they received 2.0 mL of water per day. After three days of water restriction, mice were brought to the recording room and connected to a computer system through a commutator with a 2.0-meter-long custom cable. Animals were habituated to handling and being tethered to the cable for at least 3 days. During this time, mice were connected to the computer and allowed to explore their home cage but were not exposed to the linear track. On the fourth day, behavioral recording was initiated, mice were grabbed by the tail and placed in the middle of the track and imaging was recorded within 20 seconds of the mice being introduced in the linear track. Mice were exposed to the maze every day during the learning and first 5 days following re-exposure. However, during the training period or 5 days after re-exposure, recording sessions were at different day intervals ranging from 2-5 days. The initial facing orientation of the mouse in the linear track was not controlled.

Each behavior session consisted of a 10-minute recording of the mouse exploring its home cage, without the lid and food dispenser, immediately followed (within 30 seconds) by a 20-minute recording of the mouse running on a linear track. The mouse home cage is rectangular 20 cm x 35 cm with 15 cm transparent walls while the linear track is a 1.5-meter-long track 12 cm wide with 15 cm high walls made of white plastic. The linear track was cleaned by picking up mouse droppings, spraying with 70 % ethanol, wiping the surface and waiting for at least 10 minutes before introducing another mouse. Three group of cues were place on both walls of the linear track and consisted of black stripes (2 cm wide) at different angles (figure S1d). The linear track was equipped with an automatic liquid dispensing port that would deliver between 100-200 μL of sugar water (15% sucrose in DI water) at the end of the track. The system would require the animal to run to opposite ends of the track to receive a water reward, a green LED (right side) and red LED (left side) indicated which port the mouse needed to run towards. A beeping sound was played once the mouse activated the IR sensor. Delayed reward experiments were performed by decoupling the delivery of sugar reward from IR sensor activation, thus the animal was required to wait for 5 seconds until sugar water was delivered. The beeping sound was not delayed. The position of the mouse was tracked by an ultra-wide-angle webcam at 25 Hz and 640×360 pixels positioned about 1.8 meters above the maze. Mouse position was extracted using OptiMouse (*34*).

#### Calcium imaging processing

*Video Acquisition:* signal from the microendoscope CMOS sensor was acquired thought a UVC compliant USB analog video to digital converter. The video feed was captured and saved by videoLAN media player using custom MATLAB scripts. Data acquisition was set at 25 Hz and display resolution of 720 x 576 pixels using a YUV4:2:2 codec and AVI file encapsulation. All three cameras were synchronized to star simultaneously and were verified to have latencies smaller than 10 ms. Raw videos were offline transcoded to lossless H.264 -MPEG-4 AVC codec and MP4 encapsulation and the first 2 seconds of each video were deleted. Transcoded videos were filtered using a high quality 3-dimensional low pass filter (*hqdn3d*) with spatial and temporal smoothing of 4×4 pixels and 2 frames, respectively. Denoised videos were then 4x down-sampled by a moving window averaging of 4 frames. All transcoding, smoothing, and down sampling was performed by the open source program *ffmpeg* controlled through custom MATLAB scripts. Down sampled videos were motion corrected using the recursive fast Fourier transform approach provided in the MATLAB script *sbxalign.mat* from Scanbox. For batch analysis all 4x down-sampled videos were concatenated into a larger video, motion corrected, and analyzed with CNMFe.

*Signal extraction:* motion corrected videos were saved as a matrix array and analyzed using CNMFe with a 2-fold spatial and temporal down-sampling (*22, 23*). First, we compared neuronal activity during home cage exploration and running in the linear track. All 7450 frames of calcium imaging during the 20-minute linear track task and 3750 frames during home cage exploration were analyzed simultaneously each session. In some cases, were motion artifacts were minimal, up to 200,000 frames of data was analyzed simultaneously with CNMFe and these datasets were utilized to confirm registration procedures described below. All data analysis was performed using the deconvoluted neural activity output (*neuron.S*) from CNMFe.

*Cell registration:* accurate alignment of neurons across sessions spanning days to months has noticeably become a challenging aspect of calcium imaging. Taking from studies using multiphoton imaging, registration approaches using single photon epifluorescence involves alignment of neurons between sessions based on their spatial footprint. However, spatial footprint of extracted neurons using microendoscopes can suffer from artifacts arising from changes in firing rates, extraction algorithms, and motion correction artifacts. These effects are compounded when imaging areas with dense labeling as observed in transgenic animals. To overcome these limitations, we employed an interleaved and batch data analysis approach in which a session of data was common between two days to be aligned. This common session served the purpose of decreasing noise in the extracted neuron footprints and provided firing data common to both datasets. First, the footprint extracted by CNMFe was thresholder so that only the pixels above the 50^th^ percentile formed the ROI footprint. Minor motion artifacts were corrected by a fast Fourier transform method using the PNR image output from CNMFe. The spatial correlation between all neurons with centroid distances below 15 pixels between the two alignment datasets was calculated. The correlation of the CNMFe deconvoluted neural activity was also calculated for all neuron pairs with centroid distance less than 15 pixels. Only spatial and temporal correlation above 0.5 and 0.3, respectively, were considered. An alignment coefficient was calculated by multiplying the spatial correlation and temporal correlation of every potential neuron pair. The alignment coefficient was binned in 100 intervals and the probability distribution calculated. A plot of the probability distribution showed a clear bimodal distribution (Fig. S2f), a threshold of approximately 0.5 was selected manually. All neuron pairs below this threshold were deleted, and the remaining were aligned based on an iterative selection of pairs with the highest alignment coefficient. This approach was validated by aligning a video to itself, where we observed that including temporal information increased aligning accuracy by about 10% (see supplementary information). Here, we take advantage of small motion artifacts across days in order to motion correct and analyze several days together in one animal. This ensures, that even if the neuron decreases its firing rate significantly, CNMFe will still draw a ROI around the neuron and extract its firing activity or show that it is not active. Because in some cases the FOV drifts in a non-rigid manner, we restricted our analysis to less than 30 sessions. In some figures we show the correlation image of neurons per session (ie. Figure 1f-g, and figure 3a). These images were obtained by analysis videos of CA1 calcium activity during home cage exploration and linear track (30-minute videos). The resulting correlation image from CNMFe was then motion corrected using a Fourier transform method as mentioned above.

### Data analysis

*Identifying place cells*: place fields were extracted by identifying periods when mice ran continuously faster than 3 cm/second for more than 0.4 seconds. Together, these thresholds eliminate periods during which mice were grooming, rearing, or turning. The length of the linear track was divided into bins spanning 3 centimeters (50 bins), neuronal activity at the ends of the maze (7 cm) were not included in the analysis. The average firing rate of a neuron in each bin was calculated by the sum of all calcium activity in a bin divided by the amount of time the mouse spent in that bin. The average firing rate of a neuron was then normalized by the total number of spikes in order to generate normalized tuning profile of each neuron. Neurons were classified as place cells if: (1) the place field is 15 cm wide; (2) calcium transients were present > 30 % of the time the mouse spent in the place field; and (3) the cell contains significantly greater spatial information than chance. Spatial information is calculated using (*35*):

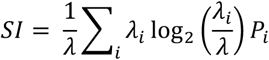

where λ is the overall average calcium transient rate of the cell, λ_i_ is the average calcium transient rate in spatial bin *i* and P_i_ is the probability the mouse is in spatial bin *i*. Chance level spatial information for each neuron is calculated by shuffling the time stamps of the calcium transients and calculating the spatial information of the shuffled trace. This is done for 1000 iterations and the spatial information of the cell is considered significant if it is higher than 95% of the shuffled traces.

*Identifying time cells*: the linear track was equipped with a LED light that would turn off once the animal activated the IR sensors at the water reward port. The ON/OFF transition of the LED in the behavior video was extracted and used as a timestamp. The LED timestamp was set as time = 0 and a window of 31 frames or the time the animal was immobile (velocity less 1 cm/sec), whichever is smaller, was used for analysis. Neurons were classified as time cells if: (1) they fire at least 20% of the times the animal activated the water port; (2) the neuron fired 30% more within its field (20 % of the time it spent immobile) than outside; and (3) the cells contain significantly greater information than chance. Temporal information was calculated using the same equation above but using λ as the average activity during immobility, λ_i_ is the average activity in frame *i* after activation of the water port. The variable Pi in this case represent the probability that the animal was not moving during frame *i*.

*Network graphs*: an adjacency matrix was generated by calculating the pairwise Pearson correlation between all neurons during a session. Only statistically significant (p<0.05) correlation values above 0.10 were used to build the adjacency matrix. The correlation threshold was also tested at 0.15 and 0.05 and similar results were observed. The weight of the edge between two neurons was set to be equal to the correlation coefficient. The networks are plotted with the network graphical tool (Gephi 0.9.2, www.gephi.com) using the built-in ForceAtlas2 layout. Modularity was calculated with Markov Diffusion approach with a diffusion time of 0.7 using the built-in function from Gephi. Cell groups are defined as cells making up each module extracted by the modularity calculation. Only the largest 11 modules were analyzed. Node metrics including degree, average clustering coefficient, eigenvector centrality, and PageRank were calculated using built in and custom MATLAB functions. The connectivity of a node in a graph was calculated by counting how many edges a node made with all other 11 modules. The module group connectivity was defined as the fraction of edges a node had with cell in the module divided by all edges that node made.

*Behavior decoding*: mouse position was decoded by a generalized linear model using the *glmfit* function with a normal distribution in MATLAB. Mouse position was discretized into 50 intervals and periods of mobility and immobility were decoded. The linear model was first trained with 50% of randomly selected place/time cell activity and mouse position and then tested on the remaining 50%. For time-lapse decoding we used the activity of neurons identified as having a place or temporal field in one session together with the mouse position to train a decoder. This decoder was then used to decode the mouse position on another day, using the current calcium activity of the place and time cells initially used to train the decoder regardless if they were classified as place or time cells.

Decoding mouse position using cell groups was performed in a similar fashion, but instead of using place/time cell activity we used the integrated deconvoluted neuronal activity of neurons making up each cell group. That is, we used the *neuron.S* output from CNMFe, identified cells in a group, binarized their deconvoluted neural activity and summed all of them across all neurons. Thus, we had 11 inputs (for each cell group) and one target (mouse position) as a training set, 70 % of the dataset was used for training. Time-lapse decoding was performed by training a decoder on one session and then testing it on session N using cell group activity as an input. Cell group activity was determined at sessions N by first calculating the maximum projection of the adjacency matrix across 5 trained sessions (5-10), calculating the cell groups using Markov diffusion. The IDs of neurons making each cell group were used to calculate their integrated group activity in all other sessions. In addition, to account for the nonlinear and asymmetric nature of fields encoded by cell groups (Fig. S10f), we also performed position decoding using a nonlinear input-output time series neural network (*timedelaynet* function in MATLAB). This network was set to have 10 hidden neurons and a delay of 360 ms (2 frames). The network was trained with 70% of randomized cell group activity (11 inputs) and mouse position, validated with 15 % and tested with 15% of the remaining data. The network trained in one session was used to decode N sessions apart. In all, cases we also tested the decoders with a randomized dataset for comparison. Error reported represent the mean absolute error ± std between the mouse position and the decoded position. All periods including learning, trained, and re-exposure were decoded. Sessions with rotated representations significantly decreased the decoders efficiency. These sessions were removed by only analyzing sessions where the mean absolute error was below the median between all sessions.

*Time/place cell decoding*: whether a neuron was place or time cell was decoded from neuronal activity by (1) calculating the intrahemispheric correlation in one session, (2) generating a graph and calculating its topological properties, (3) training a fitting neural network with half the data, (4) using the decoder trained on one session to decode another session. Graph topology metrics for each node included: degree, modularity, eigenvector centrality, PageRank, calcium transient rate, and the assemblies to which a node was connected. The calcium activity was defined as the integrated neural activity of a neuron (neuron.S matrix in CNMFe) in a session divided by the time the animal spent in the track (1200 seconds). Of these metrics, degree, module connectivity and calcium transient rate had the most effect on decoding efficiency (data not shown). All metric, excluding modularity, calcium activity, and node connectivity were normalized so that the sum in a session is equal to one. These metrics were the input to the fitting neural network and whether a neuron had (1) or not (0) a field was used as the training target. In addition, we trained a decoder simultaneously with 10 sessions of data from one mouse and used to decode which neurons were place and time cells in other animals only using their graph topology.

*Decoding stability of a cell*: we predicted whether a place/time cell we observe today will lose its field in N days by first randomly selecting 50 % of the sessions in one mouse and calculating all pairwise session intervals between them. We trained one fitting neural network for each N session interval, generating N decoders. The inputs consisted of graph topology metrics calculated from brain activity (the same parameters used above) recorded in the sessions at N intervals. The target dataset was a matrix listing whether a neuron became responsive or unresponsive to a field (logical 1) or the neuron retained its field or lack thereof (logical 0) between two sessions at a N interval. We then tested each decoder on the remaining 50 % of the sessions with intervals ranging from 1 to N. We filtered the decoder’s output so that the highest outputs above the median were considered positive.

### Quantification and statistical significance

All statistics were done with nonparametric approaches either using a Wilcoxon rank sum test, two-way ANOVA, or Friedman’s ANOVA test, or a bootstrap shuffling procedure. Correlation p-values were calculated using a two-sided t-test. Values are shown as median ± standard deviation unless stated otherwise. Two mice did not undergo the re-exposure period and were not included that analysis.

### Cell Registration across days

Cell registration across days in calcium imaging datasets obtained with microendoscopes has routinely been performed using the footprint of a neuron. However, this approach can be problematic for neurons at the periphery (where GRIN lens aberration can distort the footprint of the neuron) or in neurons with bright dendritic processes. CellReg has approached this problem by assigning scores to each registration, thus providing a useful metric to asses registration confidence (*36*). Validating registration across days using such approaches is not possible since no ground truth is available. However, the statistical nature of the CNMFe and CellReg algorithms makes it possible to analyze how minor changes in neuronal footprint can affect signal extraction and alignment. We explored the effect of analysis noise by using a 20-minute recording of CA1 activity in a mouse running in the linear track. We generated two videos, one is the original 20-minute video, and another is a temporally concatenated version of the original video (40 minute long). We motion corrected and analyzed both videos separately using CNMFe as described in the methods section. Temporal concatenation of the video increases the number of frames and changes the statistics used by CNMFe to extract neuron footprints, thus leading to minor changes in the footprints. CNMFe effectively extracted similar fraction of ROIs from both videos (649 in original vs 646 in concatenated). Next, we tested whether CellReg was able to register these ROIs using the spatial correlation of their footprint. Using a 0.5 confidence level in CellReg we found that CellReg registered 85 % of ROIs (549 out of 646) and at a confidence of 0.95 the fraction dropped to 76 % (492 out of 646). The misaligned ROIs also led to 15 % (0.5 confidence) and 24 % (0.95 confidence) increase in the number of ROIs. Thus, we observe that minor fluctuations introduced by CNMFe can lead to inaccurate registration of neurons across days when only using footprints.

Next, we investigated whether not only using footprint but also the activity profile of a neuron can enhance the registration across days. CellReg uses two metrics (spatial correlation and centroid distance) in order to build a distribution of potentially same or different cell pairs. However, these metrics are not fully independent, potentially leading to significant overlap and high uncertainty in the registration. Registration can be improved by temporally concatenating two days and setting the activity of a neuron as a constrain for alignment, a metric which is independent and orthogonal to both centroid and shape. However, this requires the ability to analyze multiple days simultaneously, which is something we can easily achieve using chronic implants thanks to the minimal drift across consecutive days **(Fig. 1b)**. To test the effectivity of this approach, we performed the same analysis as above but including in the analysis that neurons should not only look similar (0.5 spatial correlation) and be located in the same region (<15 pixels centroid distance) but should also have similar activity (0.3 temporal profile). Using this method, we can align 93 % of all the ROIs (599 out of 646 ROIs), leading to 7% increase in the number of total ROIs. However, not all ROIs generated by CNMFe can be considered neurons with certainty, some may be segmented dendrites or background fluctuations (Fig. S2b-c). We removed these ROIs from the pool of registered ROIs by selecting only those with particular area (30 to 250 pixel^2^), circularity (less than 4.0), and PNR (above 8). There were 80 ROIs not satisfying these constrains in the original video and 66 in the concatenated version. After removal of these ROIs, the fraction aligned using our method (90 %) or CellReg (82 %) did not change dramatically. Using our method, we calculated that analysis of two versions of the same data by CNMFe resulted in footprints with areas differing by as much as 19 ± 18 pixel^2^.

We tested whether the improvement persisted when analyzing real data. In this case, we analyzed three 20-minute recordings of CA1 activity in the linear track separated by 1 day **(Fig. S2a)**. Two sets of recordings were generated. Set A contains temporally concatenated recordings of day 1 and day 2 (in that order) and set C contains recordings from day 2 and 3 **(Fig. S2d).** The recording of day 2 is common to both sets and the calcium activity of such neurons can be assigned as a constrain on the alignment. That is, registered neurons should not only look similar but should fire equally on both sets. We also concatenated all three videos into a 60-minute video (Set D) and analyzed it with CNMFe. Set D can be used as validation, in this set we observe that most (97 %) ROIs are active in all 3 days and there is a total of 731 ROIs **(Fig S2d)**. CNMFe extracted 702 ROIs in Set A and 700 ROIs in Set C. CellReg is able to align 74 % of these ROIs at 0.5 confidence (521 out of 700) and 63 % at 0.95 confidence (442 out of 700). Due to the misalignment, a total of 881 ROIs are detected in both sets, an error of 21 % when compared to the 731 ROIs detected by CNMFe. Using our method, we can align 87 % of the ROIs (607 out of 700) and identify a total of 794 ROIs (13 % error compared to 731 ROIs by CNMFe). CNMFe found that 97 % of ROIs are active all three days, our method identifies that 88 %, and CellReg found 72 %. All three approaches lead to the same answer; most neurons are active all three days and similar results were obtained when extending the analysis to longer timescales. Due to the more reliable registration obtained by batch analysis using CNMFe as well as the higher detection levels **(Fig. S2b-c)**, this method was used in this study. However, analyzing the data using the temporal constrain mentioned above provided identical results.

## Supplementary Figures

**Supplementary figure S1.**
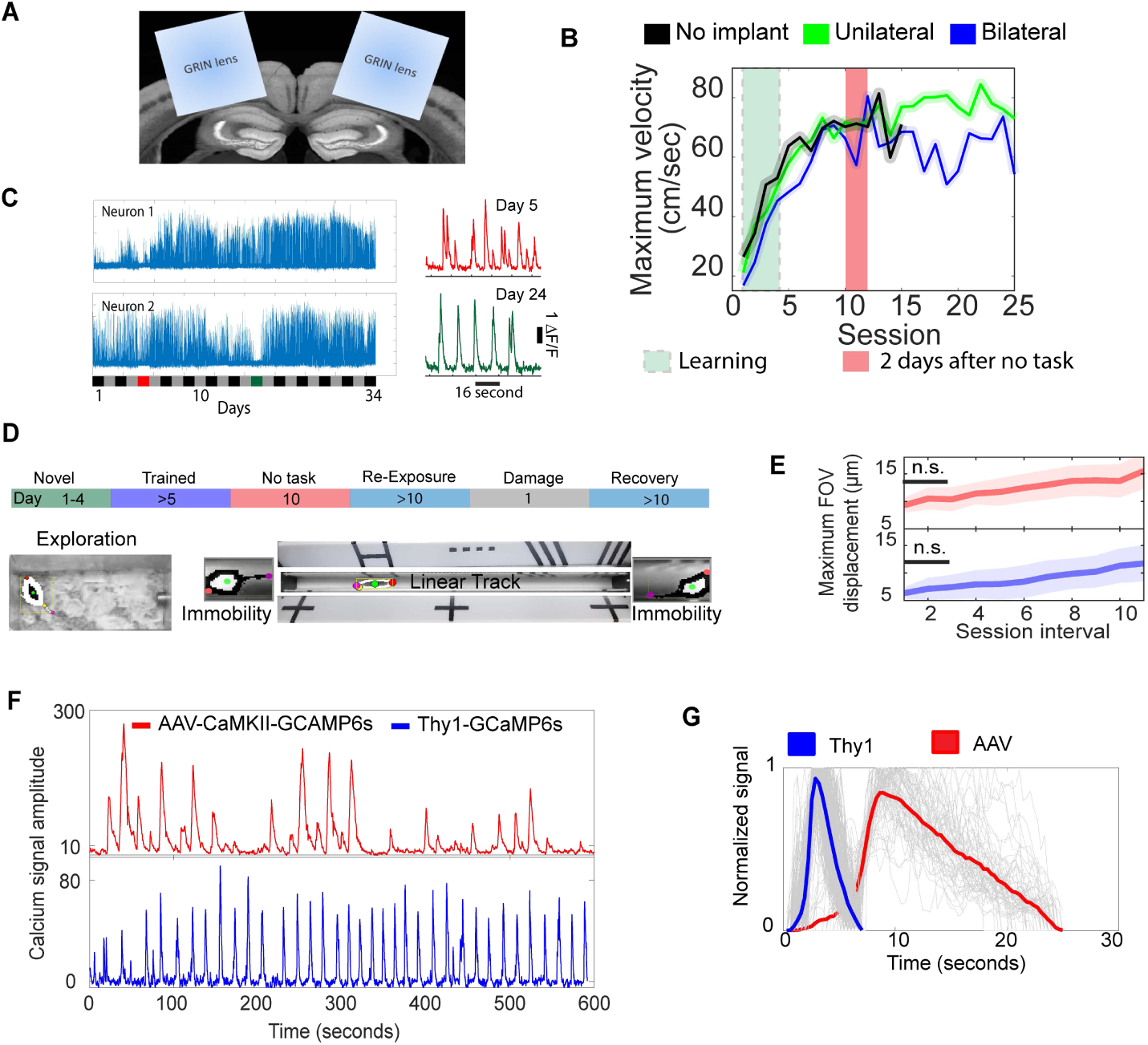
Simultaneous bilateral imaging of CA1 activity in freely moving animals. **(A)** Coronal section of a mouse brain with diagram of bilateral 1.8 mm GRIN lens implants. **(B)** Bilateral and unilateral implants do not affect the maximum velocity in the linear track. Rectangles highlight period of novelty (first 4 sessions) and initial 2 days following re-exposure to the task (n = 3 no implant, n = 4 bilateral, n = 5 unilateral). Also, the mouse behavior does not degrade if the animals are not exposed to the task (red rectangle. **(C)** (left) background corrected calcium signal from two neurons showing continuous activity throughout several days. (right) Inset showing several seconds of activity on two days of neuron 2. **(D)** Experimental schedule and tracking of mice in the home cage (left) and linear track (right) during periods of running and immobility (nose, tail, and center of mass shown in red, pink, and green). Cues shown on top and bottom of the linear track. **(E)** Motion artifacts across 3 days are not significantly different than same-day fluctuation (p<0.05, 3 sessions apart drift 9 ± 5 μm, 10 sessions drift 15 ± 5 μm, lines represent median ± std, ranksum-test). **(F)** Calcium transients in CA1 neurons of a mouse running in a linear track using adeno associated viruses to induce expression of GCaMP6s (red trace) and using transgenesis to induce expression (blue traces). AAV data was obtained from www.miniscope.org. **(G)** Interestingly, the fastest decaying calcium transients we observed had a half-life of 0.56 ± 0.14 s (mean ± s.d., n=554), about 5-fold faster than that observed using AAV GCaMP6s, (2.9 ± 0.5 s, n = 63, p<10^−32^, rank-sum test). This is similar to the faster dynamics observed in neuron L2/L3 neurons of the anterior lateral motor cortex of transgenic animals compared to viral vector mediated GCaMP6s expression (*19*).

**Supplementary figure S2.**
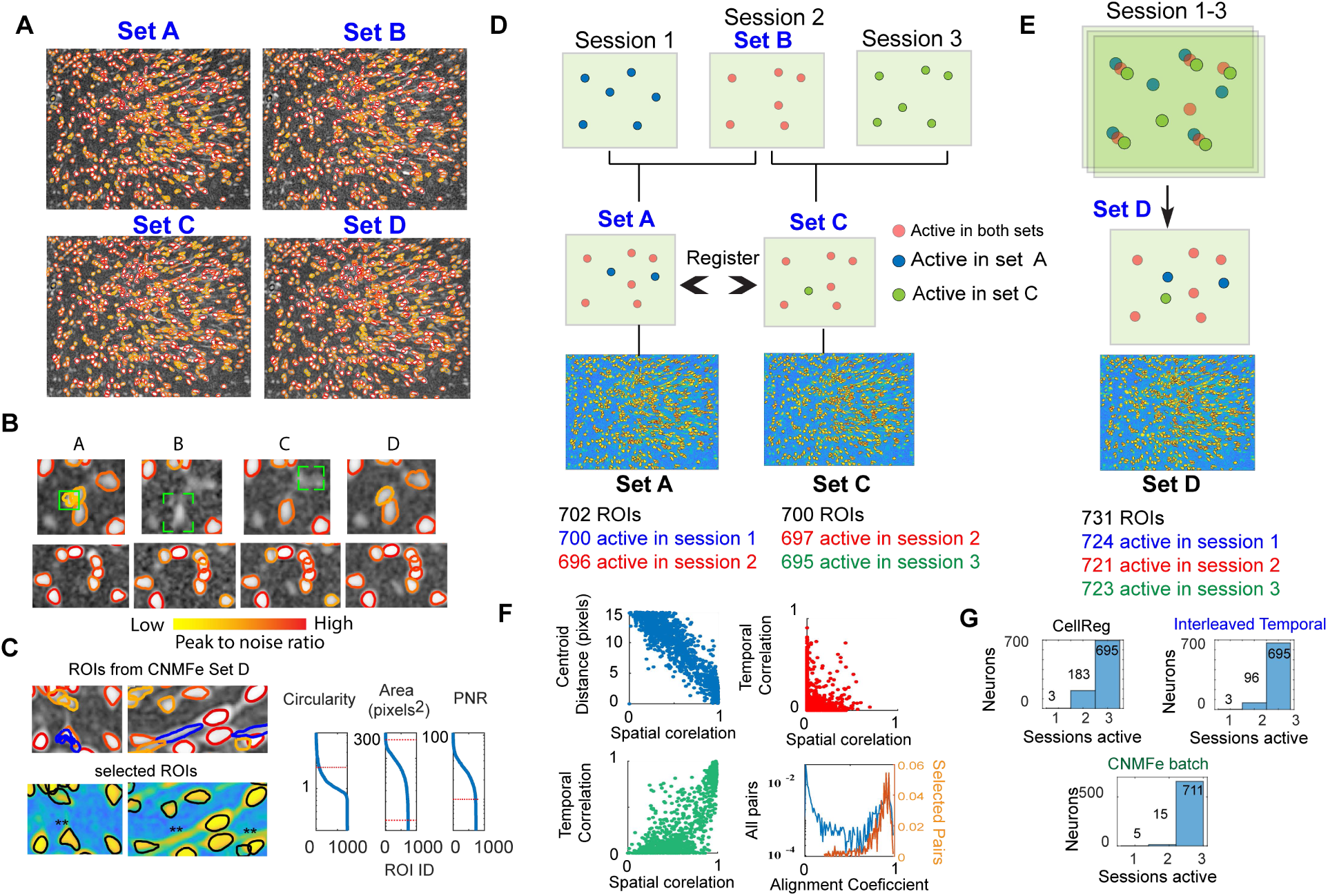
Using temporally concatenated datasets improves cell registration across days. **(A)** Three 20-minute recording of CA1 activity in a mouse running in the linear track on three different days were combined in into 4 sets. Set A includes day 1 + 2 (14900 frames), Set B is only day 2 (7450 frames), Set C includes days 2 + 3 (14900 frames), and Set D contains days 1 + 2 + 3 (22350 frames). **(B)** Concatenation leads to better signal extraction using CNMFe. Set D performs better than all other sets. Note the regions highlighted by the green rectangle where there is some signal but not enough to be identified as a neuron. The lower panels show another region four very close neuron are better extracted by combining all videos. **(C)** Two small regions from set D identifying dendrites (right, blue trace) and residual noise (left, blue trace) as neurons. These ROIs were removed from the analysis by setting shape and peak-to-noise ration thresholds (right panel). **(D-E)** Diagram showing concatenation approach and CNMFe output. Active neurons were determined from the deconvoluted neural activity (neuron.S) output from CNMFe. **(F)** Steps used in the interleaved approach tested here. (top left) Relationship between centroid distance and spatial correlation of all ROIS with centroids less than 15 pixels away. (Top right) Temporal and spatial correlation of ROIs with centroids further than 15 pixels. (Bottom left) Temporal and spatial correlation of ROIs with centroids closer than 15 pixels. (Bottom right) Alignment coefficient of all ROI pairs (blue) and those registered across Set A and Set C (orange trace). Note the better separation of distributions in the lower left panel compared to the top left panel. **(G)** Results from all three approaches indicating that most neurons are active most days.

**Supplementary figure S3.**
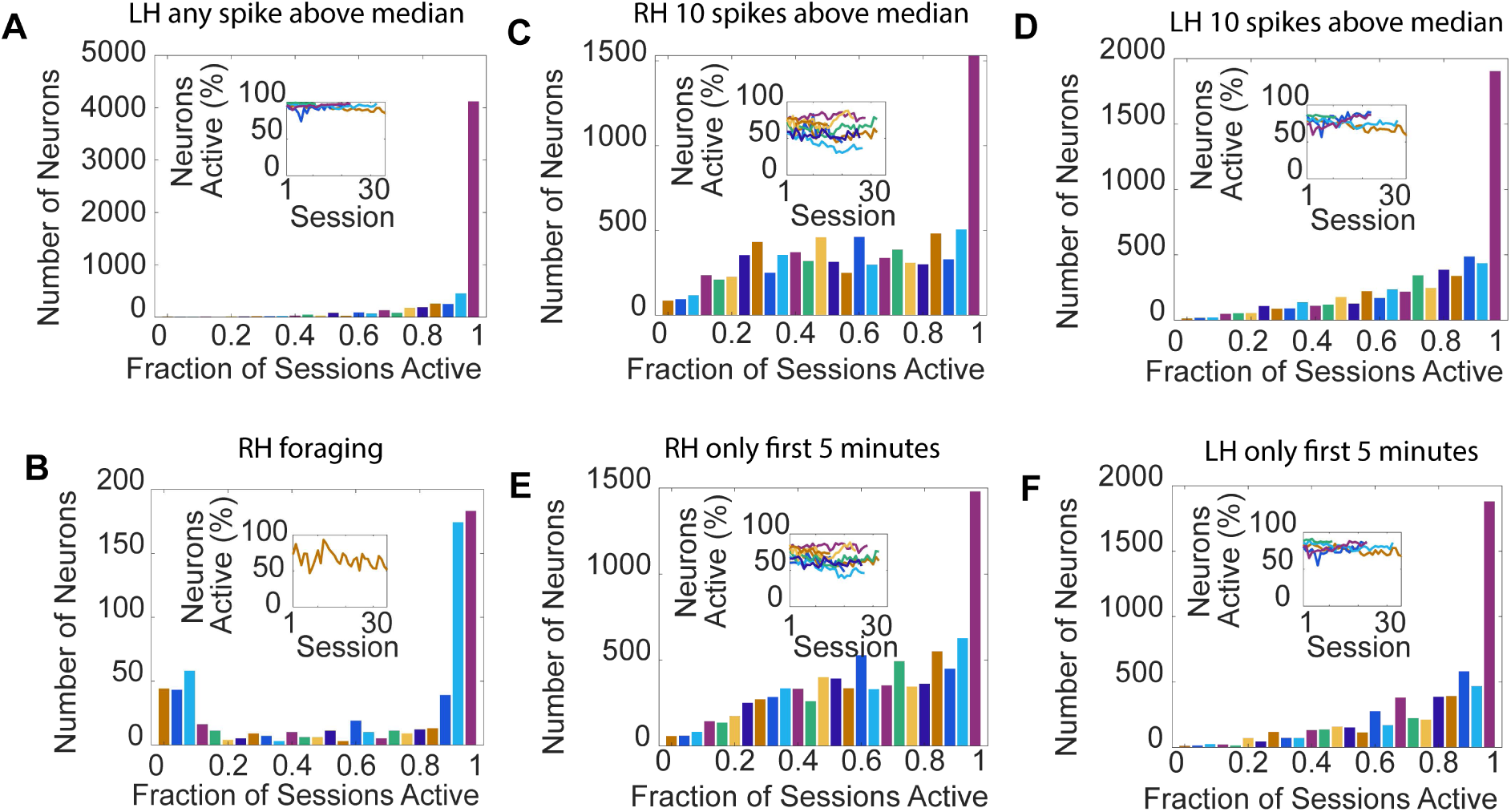
Neurons are active most days. **(A)** Cumulative activity distribution of the left hemisphere showing that the majority of neurons are active on most sessions (n=5 mice). Inset shows fraction active on each session for each animal. A neuron is classified as active if it had a calcium transient above the mean calcium signal across all neurons during that session. **(B)** Same as (a) but during a 10-minute recording while the mouse explored its home cage. **(C-D)** Only considering a neuron is active if 10 calcium transients were above the mean calcium signal in a session (linear track data only). **(E-F)** Only considering the first 5 minutes of recording in the track, active defined as panel a.

**Supplementary figure S4.**
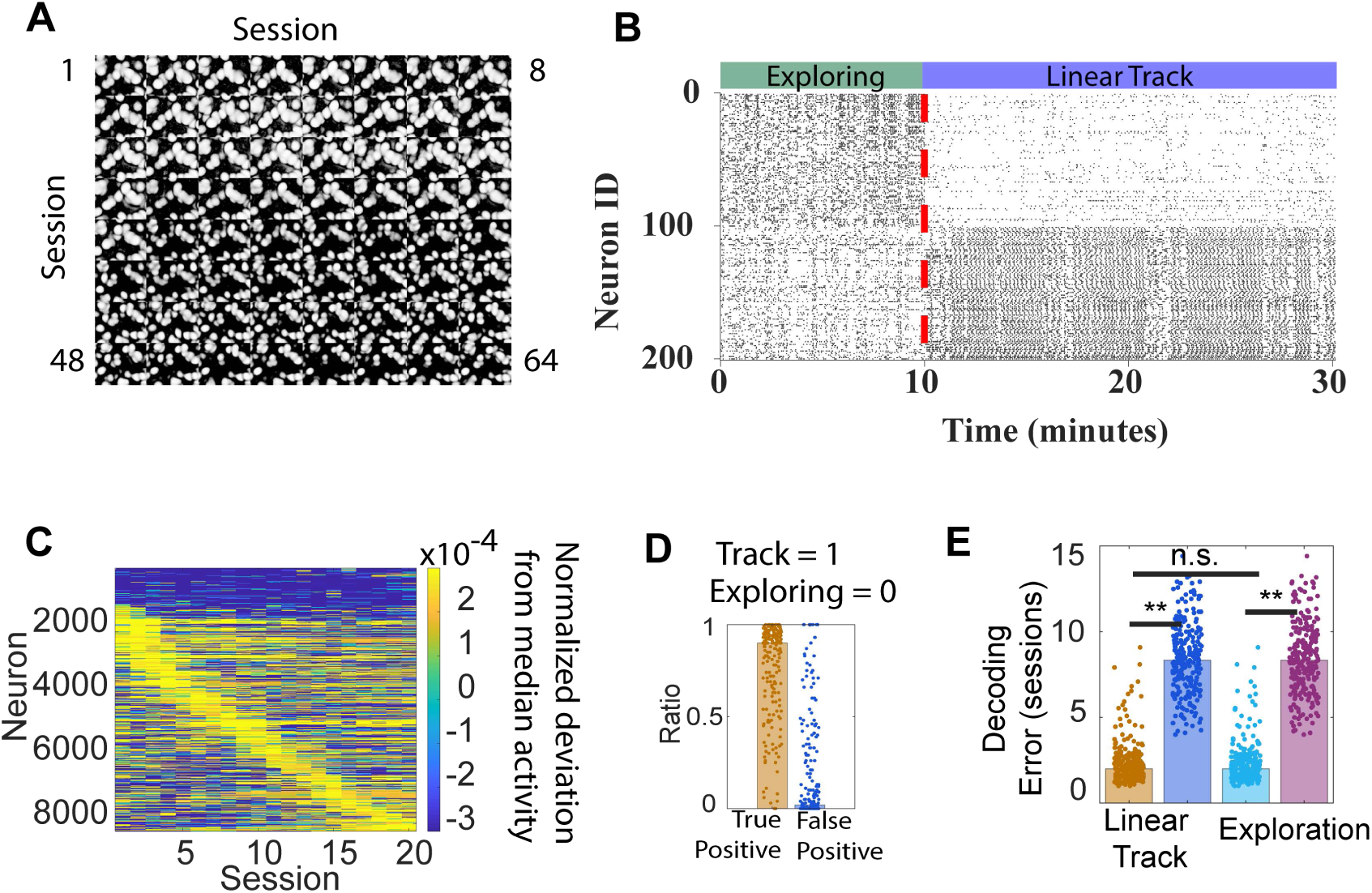
Neurons change their firing rate across environments and days. **(A)** Correlated pixel intensity image of the same region shown in Fig. 1f but across 64 sessions spanning 8 months. There is a 2-month interval between session 33 and 34. Note the gradual change in ROIs due to small drifts in the focal plane of illumination. **(B)** Despite their stable participation, neurons change their firing rate across environments and days. Raster plot of deconvoluted activity from 100 neurons with increased activity during exploration (ID: 1-100) and while running in the linear track (ID: 101-200). Vertical line shows the transition from one environment to another. **(C)** Firing rates change between days. Normalized burst deviation from the population median for all mice across days. For each row we calculated how many bursts were above or below the median population burst for that session and then normalized so that the sum of all burst deviations equals one. Neurons were sorted by activity deviation. **(D)** Accuracy of a fitting neural network trained with the neuronal activity of 200 neurons with preference for the linear track or home cage (panel b) to identify whether the animal was in the linear track (positive) or in the home cage (negative). Average true-positive-rate 0.86 and false-positive 0.11 (p<10^−10^, each dot represents a session). **(E)** Bayesian classifier trained with neuronal rate changes (panel c) in the right hemisphere to identify the environment and session, respectively. Average decoding error for the track (orange) and home cage (cyan) were similar (1.8 ± 0.5 and 1.7 ± 0.5). The blue bar and violet show the error if the inputs (firing rates, rows in panel c) are randomized.

**Supplementary figure S5.**
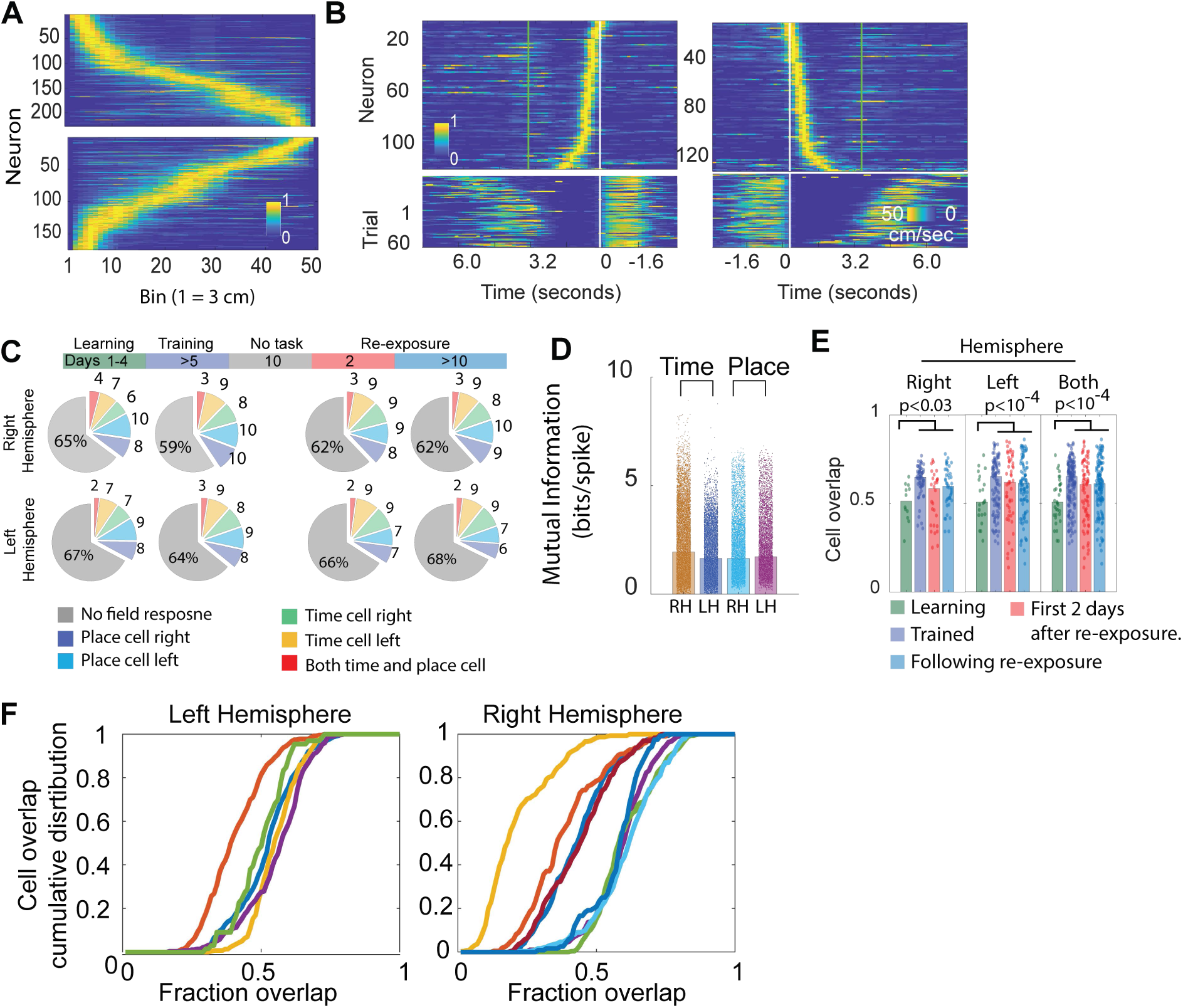
Custom microendoscopes in transgenic mice can detect neurons with spatial and temporal fields. **(A)** Normalized tuning profiles of neurons with statistically significant place field when running right (top) and when running left (bottom), only data from the right hemisphere in one mouse during one session is shown. **(B)** Normalized tuning curves of time cells firing at the left side (left) and right side (right) of the track after activation of the sugar reward port shown in white lines (the white-green lines indicate the cutoff window for calculation of time fields). (Bottom) Mouse velocity on each running trial aligned to the time after reward delivery (white line). **(C)** During periods of mobility and immobility we observe similar fractions of neurons with statistically significant fields (15 ± 7 % vs 12 ± 6 %, p>0.7, n = 12). Neurons active during immobility had partial overlap with place cells (14 ± 9 %), were slightly more direction selective than place cells (% bidirectional, 4 ± 3 vs 6 ± 5, p<10^−14^, n =313 sessions), and failed to fire within their field more often than place cells (% burst within field, 65 ± 13 % vs 79 ± 6 %, p<10^−5^, n=12). **(D)** Mutual information of place and time cells in all mice across all sessions (each dot is one cell). The mutual information of place and time cells was similar (in bits/spike, 1.8 ± 1.1 for time cells and 1.7 ± 1.3 for place cells, p>0.5, Kolmogorov-Smirnov on distributions). **(E)** Cell overlap of place and time cells during learning (first 4 sessions), trained (sessions 5 to before the no-task), the first 2 days following re-exposure (red) and the subsequent days (ranksum test, each do represent as session pair). **(F)** Cumulative distribution of cell overlaps for all in individual mice in each hemisphere.

**Figure S6.**
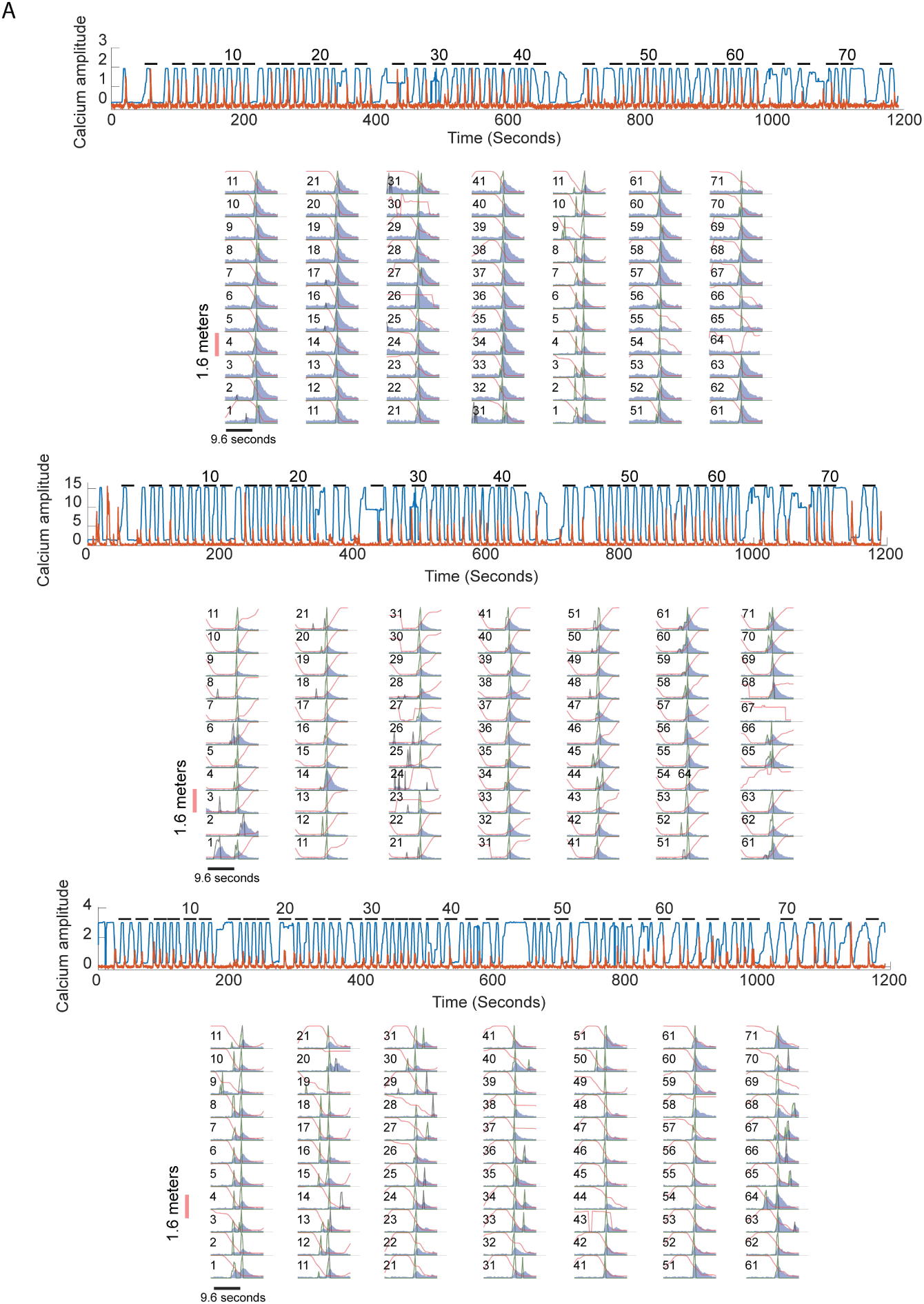
Three place cells in the linear track showing robust response to their receptive fields. The top panels show the complete 20-minute session in the linear track and the bottom panels a ∼ 20 second window near the maximum activity of the cell. The blue trace represents the background subtracted calcium transient and it is normalized so that the maximum during the session is equal to one. The black and green lines represent the spikes included and not included in the analysis, depending on whether they occur while the animal was moving or not, respectively. The red line shows the position of the mouse in the track. Numbers indicate the trial number and every even trial is shown on the top panel.

**Supplementary figure S7.**
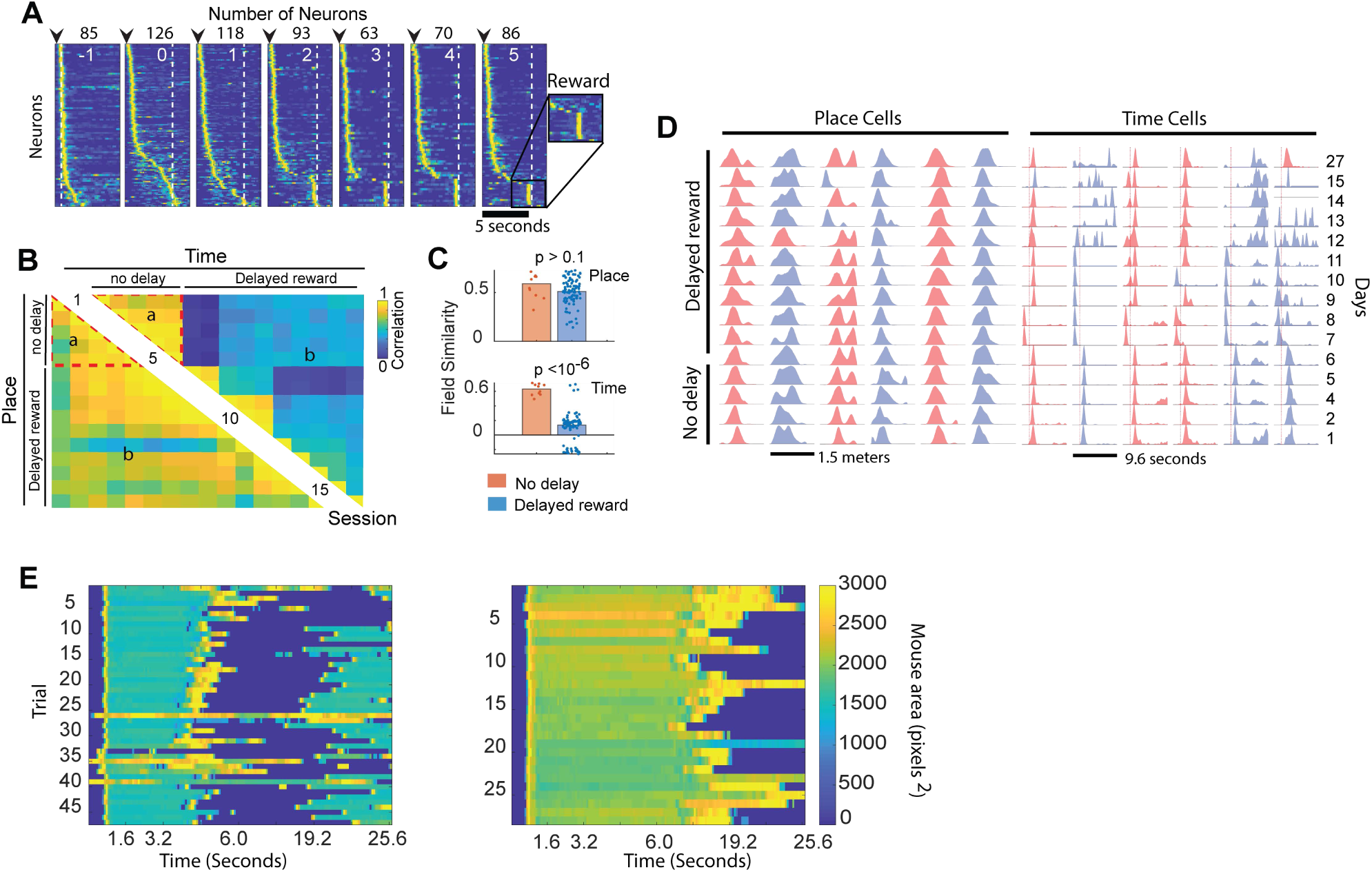
Changes in reward time affects the field response of time but not place cells. **(A)** Tuning curves of time cells at the right side of the maze during a delayed reward period. White numbers indicate sessions after delay and the black numbers the number cells identified in the session. Black arrows indicate activation of the IR sensor and the dashed vertical line shows the time of water delivery. Neurons are sorted by the delay between their maximum activity and activation of the water port. Neurons are not aligned, and each row may correspond to different neurons. **(B)** The field similarity of time cells changes after the delayed reward period, but place fields remain stable. Pairwise field correlation of neurons which retained their fields between pairs of sessions. Colors represent the median value of a session pair. Numbers indicate the session. Sessions were grouped as indicated by the dashed red boundaries, letters are labels for each region. **(C)** Quantification of field similarity of place cells and time cells before and after introducing a 5 second delay to delivery of sugar water (ranksum test, each dot represents median correlation of a session pair). **(D)** Normalized response fields of six place cells (left) and time cells (right) during the delayed reward period. **(E)** To quantify whether mice moved or not during the delayed reward period we used the mouse area extracted with OptiMouse. Normally we observed that mice did not move for about 4 seconds after activating the water reward port (left); however, mice did not move for about 15 seconds when the reward was delayed by 5 second (right).

**Supplementary figure S8.**
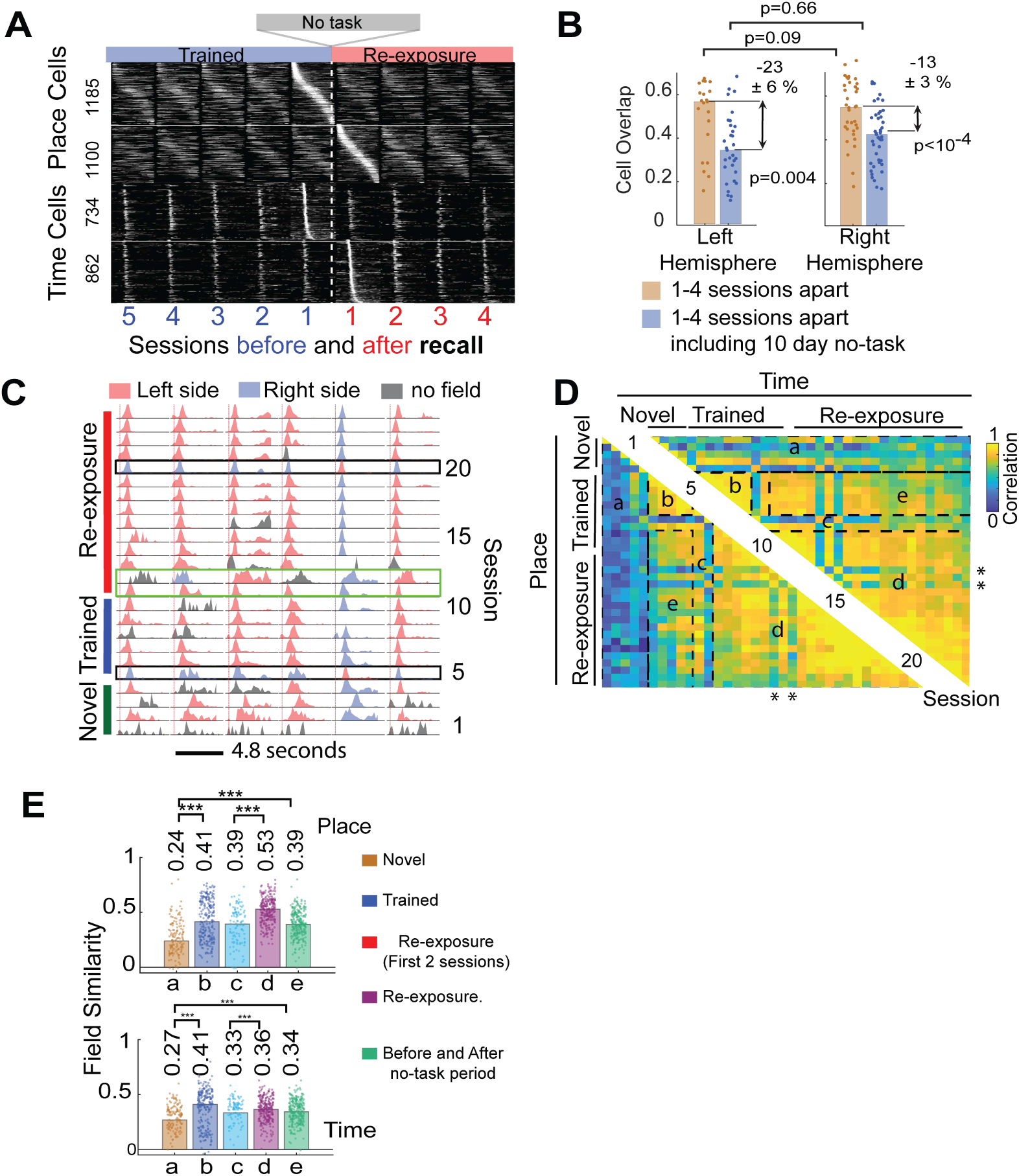
Place and time field persist through periods of no exposure to the linear track. **(A)** Normalized tuning curves of all place cells (top) and time cells (bottom) on the right hemisphere aligned to the day before and after the no-task period. **(B)** After re-exposure to the linear track, 13-23 % of place/time cells became unresponsive to their field. This figure is analyzed differently than figure 2d in order to get a wider range of comparison between session intervals. This approach was taken to minimize the potential effect of re-exposure on cell overlap and to be able to analyze each hemisphere individually. In this case we calculate the cell overlap between all possible intervals between sessions 1-4 before the gap (shown in blue numbers in panel a). Then we compare what the cell overlap is between all possible intervals between the session before the gap (blue “1”) and sessions 1, 2, 3 after the gap shown in red (median ± sem, ranksum test). **(C)** Response fields of six cells with fields during periods of immobility in the right hemisphere across 45 days. Black rectangles show sessions in which 88 ± 8 % of fields changed direction. Green rectangle are the two sessions after re-exposure. Red vertical line in (C) indicates activation of the water reward port. **(D)** Pair-wise field correlation of neurons in the left hemisphere with a field response between sessions. Colors represent the median value. Numbers indicate the session. Sessions were grouped as indicated by the dashed rectangle and letters. This is a different animal than Fig. 2f and is shown to highlight period of rotations happening during re-exposure (region d, marked with asterisk). **(E)** The similarity between fields in each group of sessions was obtained by subtracting the correlation obtained by random aligning neurons (n = 5, left hemisphere, each dot is the median between two sessions, p< 10^−4^ marked by ***). These calculations include sessions with rotated fields.

**Supplementary figure S9.**
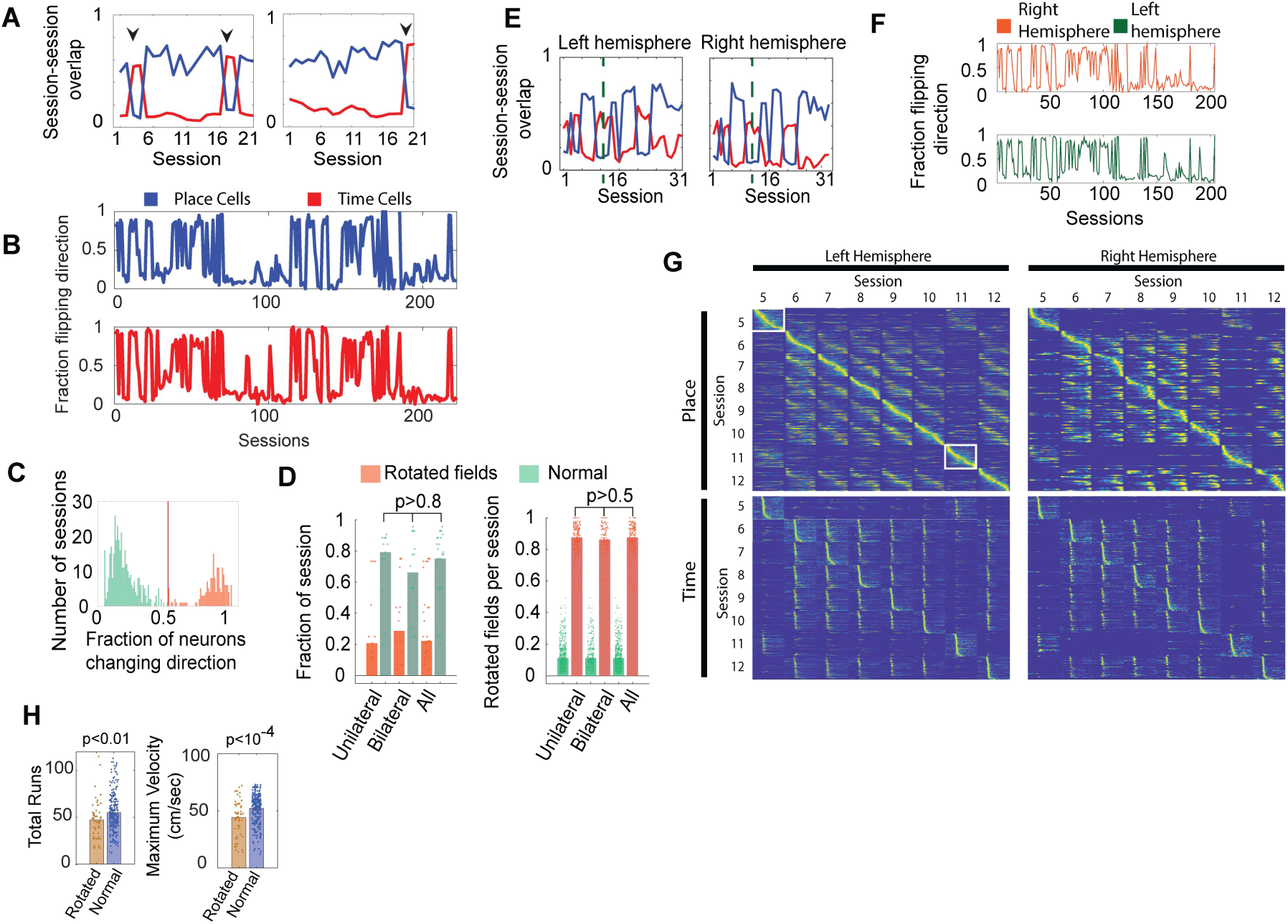
Place and time representations undergo spontaneous global rotations. **(A)** Direction specific overlap of place and time cells between two consecutive sessions in the right hemisphere. The blue trace represents the overlap between same direction neurons in session N and N+1. The red trace represents the overlap of neuron with one direction in session N with neurons of the opposite direction on session N+1. An increase in red and decrease in blue traces indicates change in directionality of place/time cells, shown with black arrows. The left and right panel is calculated from two animals being exposed to the linear track on the same days. Rotation events are not correlated across animals, indicating that these changes are not due to major changes in the environment. **(B)** Place cells (blue trace) and time cells (red trace) change their direction preference simultaneously. Plot shows the fraction of place or time fields rotated per session (n = 222 sessions, 4 bilateral mice, correlation 0.83, p<10^−10^). **(C)** Distribution of session with rotated place and time representations (orange) and without rotated representations (green), red line indicates threshold used to identify rotated sessions (n=12, place and time counted separately). Approximately ∼30% of all sessions were rotated (110 rotated, 255 normal, n=365 sessions in 12 mice, both hemispheres). Between normal sessions we observe 11 ± 9 % of place/time cell change their field; however, in rotated sessions 87 ± 8 % change. **(D)** The fraction of sessions with rotated representations was the same for unilateral and bilateral animals (0.21 ± 0.26 unilateral, n = 5, 0.27 ± 0.27, bilateral, n=4, and 0.22 ± 0.26 % for both, ranksum test). Time and place cells of each hemisphere analyzed independently. The fraction of neurons undergoing rotations was also the same. **(E)** Same as (a) but in a mouse with bilateral implants showing that both place and time cells in both hemispheres change direction simultaneously. Dashed vertical line highlight a 10-day period of no task. **(F)** Fraction of place and time cells in four bilateral mice highlight simultaneous switching across hemispheres (correlation 0.85, p<10^−10^). **(G)** Normalized tuning profiles of place and time cells across sessions in one mouse showing that field rotations happen in both hemispheres. **(H)** During sessions with rotated representations mice ran less distance (total number of laps) and they reached a lower maximum velocity (ranksum-test, each dot represents a session).

**Supplementary figure S10.**
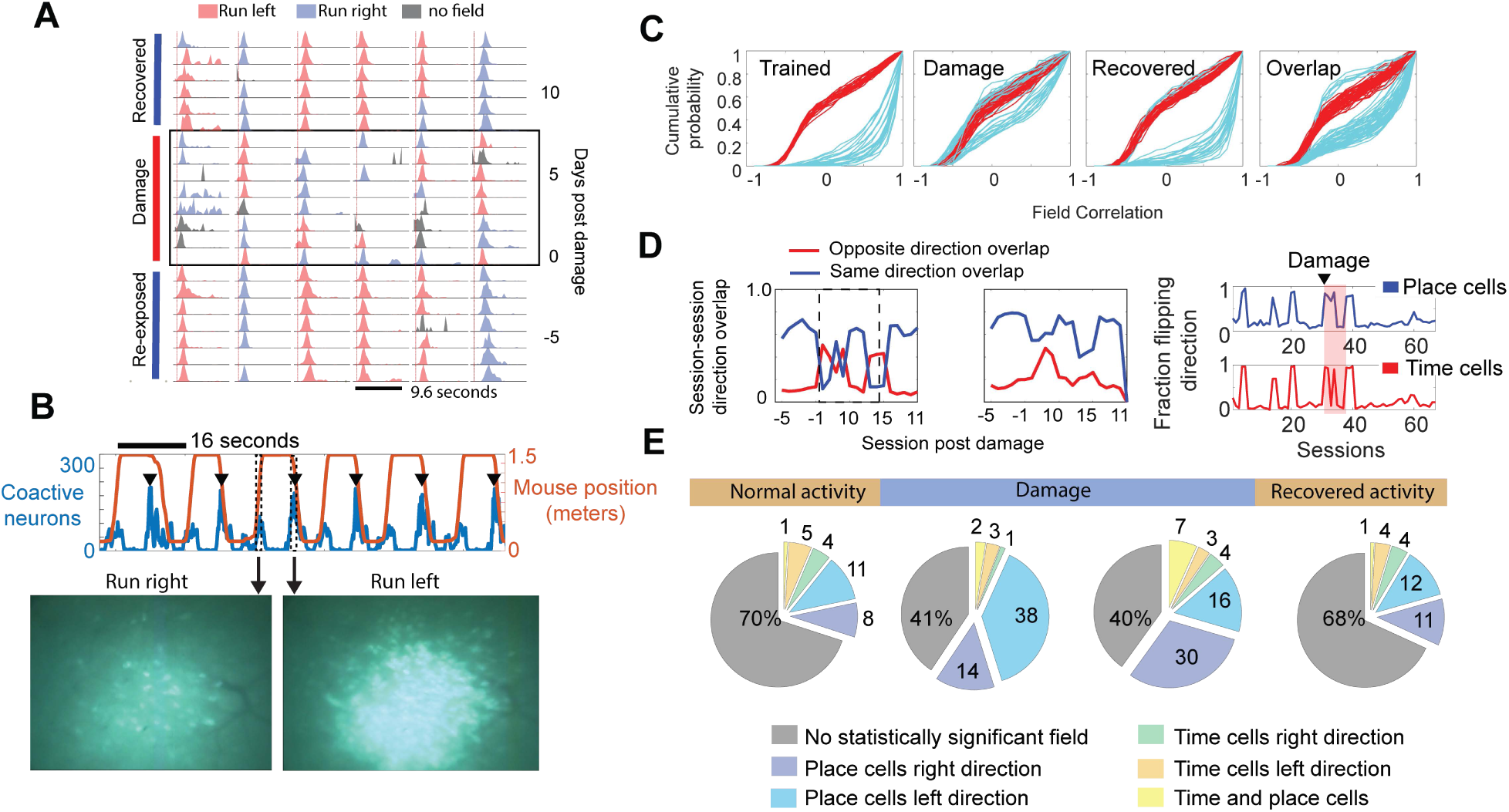
High illumination damage induces place specific abnormal activity, increase in place cells, and increase rotation events. **(A)** Normalized response fields of six place time (right) cells before, during, and after the lesion in CA1. **(B)** (top) Blue trace represents the number of coactive neurons during a frame (160 milliseconds), reaching approximately ∼250 neurons (black arrows) in one direction but only ∼ 100 in the other. Orange trace shows the position of the mouse in the linear track. (Bottom) Single frame capture during a normal running right period and a burst while running left. **(C)** Cumulative distribution of pairwise field correlation between place cells during learning, trained, and recovered periods. Red traces represent the field correlation of random place cell pairs between two sessions. Cyan trace represents the correlation of place cells that retained their responses across two sessions. The overlap traces represent the similarity between fields before and after the damage. **(D)** Following damage to CA1 the incidence of place and time field rotation increased in this animal (4 out of 7 sessions, left panel) compared to before and after the abnormal activity (8 out of 60 sessions, right panel). The right panel represents sessions recorded across 7 months and analyzed in 3 datasets of 21-25 sessions each. **(E)** The fraction of neurons with statistically significant place field also increased during abnormal activity (normal 18 ± 10 % compared to 43 ± 13 %, p<10^−4^, ranksum test).

**Figure S11.**
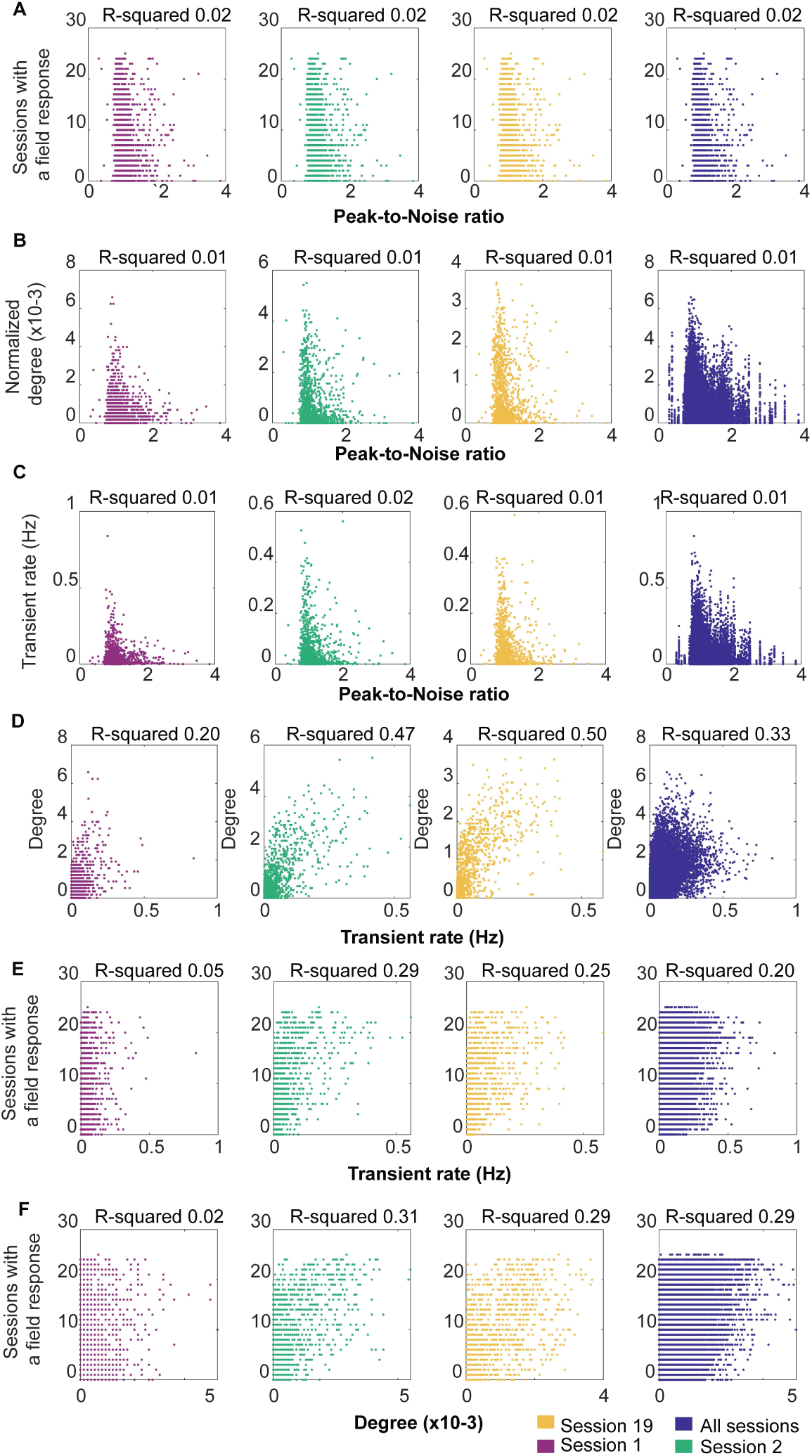
Place/Time field stability are not due to detection problems. We observe a near exponential distribution of peak-to-noise ratios (PNR) (**Fig. S2c)**. However, there is no relationship between **(A)** the number of days a neuron was classified as having a field response and PNR (p<10^−10^). **(B-C)** The degree of a neuron in a graph or the firing rate had also no dependence on PNR (p<10^−10^). **(D)** As a population, there was a small relationship between the degree of a neuron and its firing rate (R-squared 0.33, p<10^−10^). **(E)** The relationship between the days a neuron was responsive to a field and the firing rate was also small (R-squared 0.20, p<10^−10^). **(F)** The number of days a neuron was classified as having a field response was slightly related to the degree of the node representing that neuron in the graph (R-squared 0.29, p<10^−50^). Note that figure S11B and S11E is calculated using all neurons, whereas figure 4J is calculated using only place and time cells.

**Supplementary figure S12.**
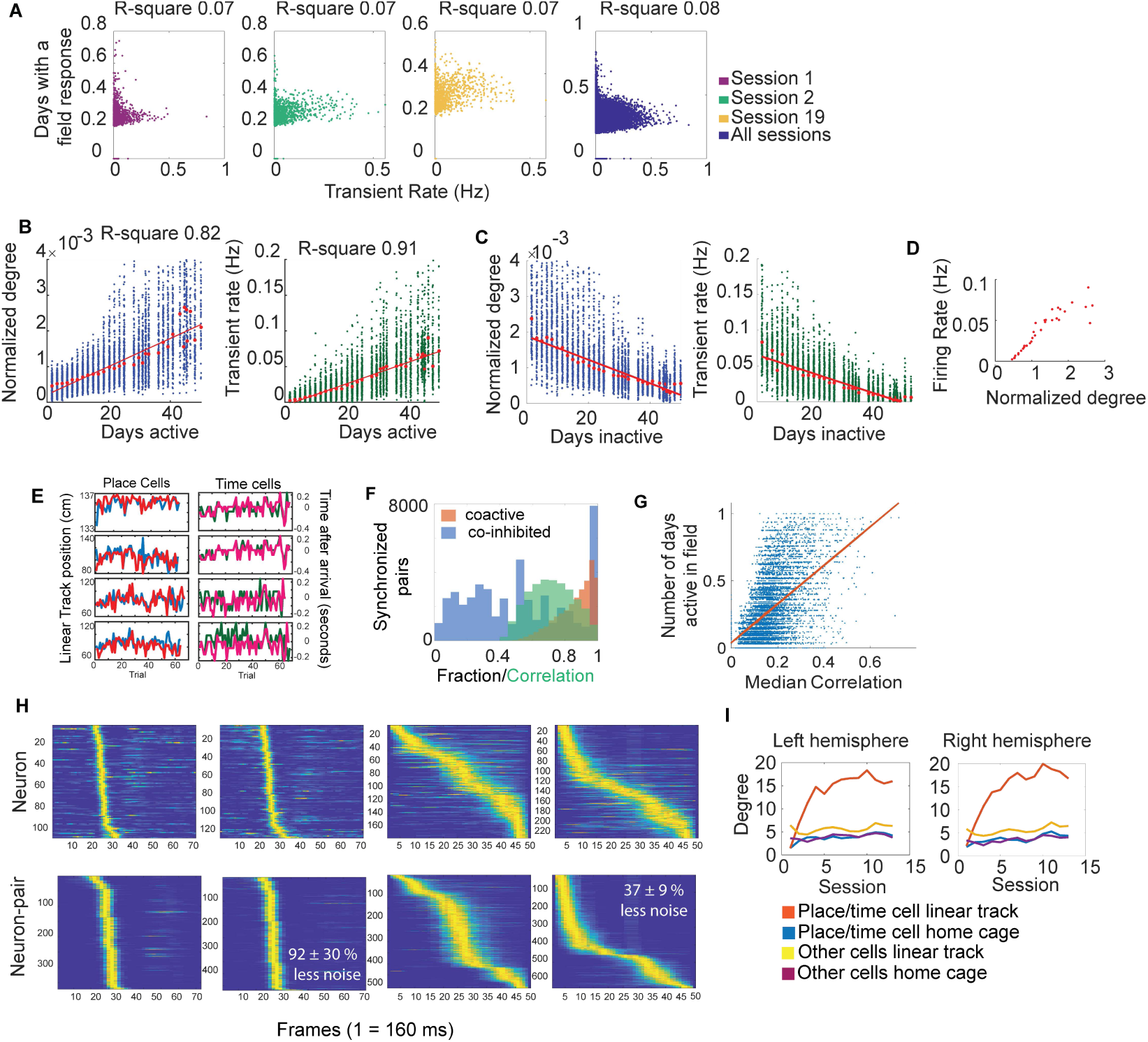
Degree is proportional to calcium transient rate but provides more information. **(A)** We observe that the Pearson’s correlation between two neurons is independent of the transient rate of each neuron in the pair (how many deconvoluted calcium events per seconds). Indicating that more active neurons do not have higher correlation. **(B)** Even though we observe a minor relationship between how many neuron pair a neuron has (degree) and its calcium transient rate (Figure S11d). The relationship between degree or calcium transient rate and the number of days a neuron has a field response becomes significant when only considering place/time cells. Indicating that more active cells have more neuron pairs and are more stable. **(C)** There is also an inverse relationship between degree or calcium transient rate and whether the neuron had spatial or temporal information. Neurons which were never classified as having a field response had the lowest degree/transient rate. **(D)** The relationship between degree and transient rate become significant when only considering place/time cells (n=12 mice, median, R-squared 0.91, p<10^−10^). **(E)** The correlation between neurons is not by chance since neuron pairs deviate from their field centroid Four neuron pairs of place and time cells shown. Time cell field ‘position’ are measured in seconds after or before activation of the reward port. **(F)** Neuron pairs are co-active 90 ± 3 % (orange distribution) of the time and when one does not fire in its field, its neuron pair does not fire 50 ± 20 % of the time (blue distribution). Synchronized pairs also show correlated fluctuations of their fields (green distribution, 0.77 ± 0.17, p<10^−8^). **(G)** The number of sessions neuron pairs remain responsive to a field is proportional to the level of synchrony in the neuron pair (R-square 0.35, p< 10^−4^). **(H)** Neuron pairs encode spatial and temporal information with less noise than transient rate alone. Time cell pairs 4.6±. 1.1 bits/spike vs individual neurons 2.3 ±. 0.5 bits/spike, p<10^−7^, ranksum test. Place cell pairs 2.1 ±. 0.3 bits/ spike vs individual neurons 1.5 ± 0.2 bits/spike, p<10^−7^, ranksum test. For these calculations we deleted every spike that happened out of sync in neuron pairs with significant correlation above 0.2. Then we used the noise free (out of sync signal removed) neuronal activity to calculate place and time fields as described in the methods. (I) Place and time cells become synchronized with multiple pairs in the linear track but not while exploring in the home cage.

**Supplementary figure S13.**
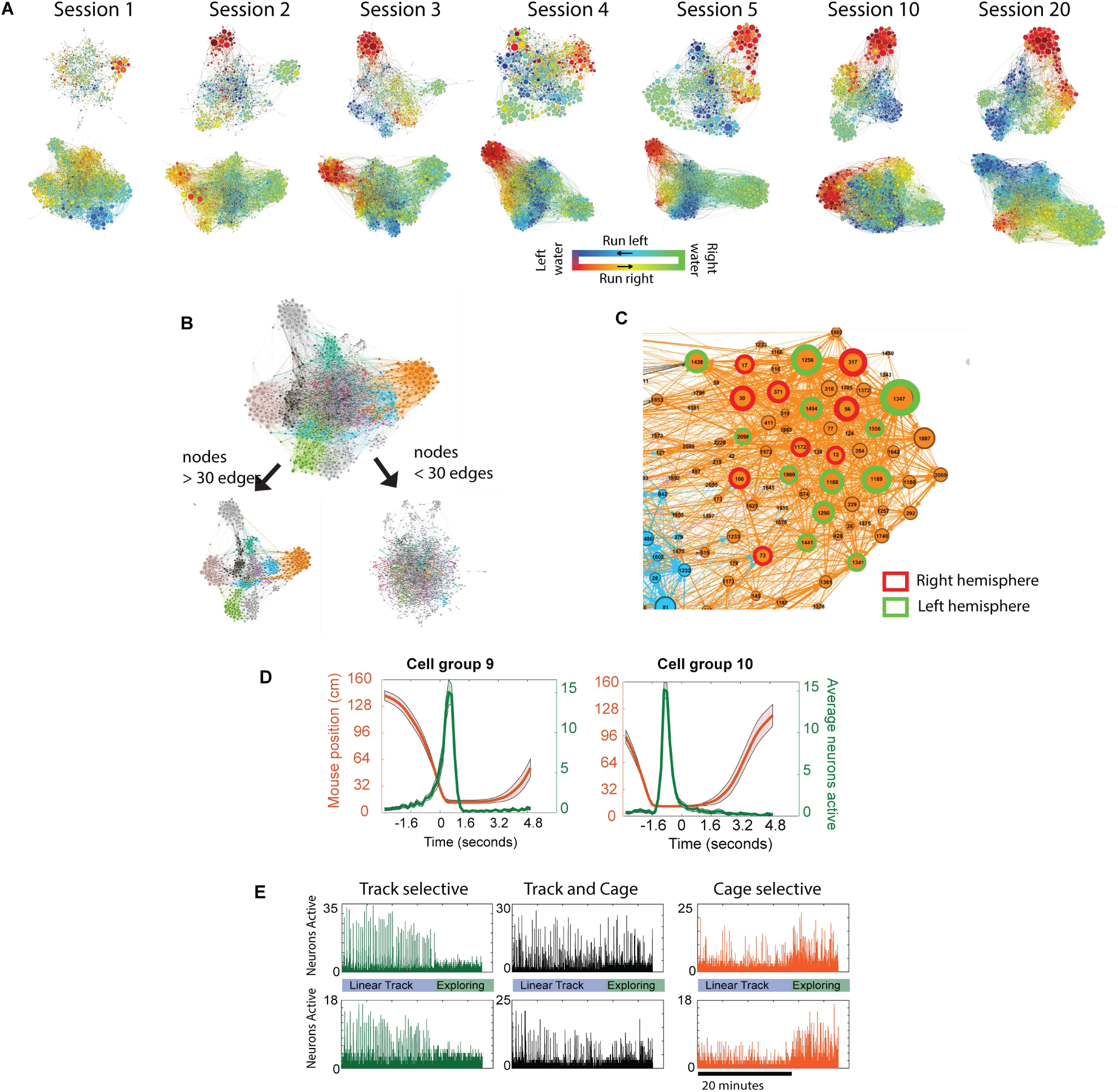
Intra- and interhemispheric synchrony in CA1 develop during learning and encodes information about the task. (A) Network graph of two mice during the learning phase (first 5 sessions) and after the mice have become habituated to the task (sessions 10 and 20). Note that opposite ends of the linear track (green vs red/blue) are separated by a largest distance, indicating that neurons on either of these ends have little correlated activity, as expected. **(B)** Network graph of neuronal activity in both hemispheres of a mouse running in the linear track. Colors indicate cell groups identified using the Markov diffusion approach. The graph can be decomposed into a graph with small world topology and a random graph by selecting nodes with more than or less than 30 edges, respectively. Thus, highly synchronized neurons form part of a functional network with small world topology spanning both hemispheres. **(C)** Close in view of the cell group in (c) with a subset of neurons from the left hemisphere (green) and right hemisphere (red) shown. **(D)** Two cell groups with place (left) and time (right) preference. Note the asymmetric shape of the distributions, indicating that as a group, neurons could use non-linear activity in order to encode position. **(E)** Two cell groups with robust synchronous activity in the linear track but not the home cage (green), two cell groups with synchronous activity both in the home cage and linear track (black), and two cell groups with synchronous activity in the home cage.

**Supplementary figure S14.**
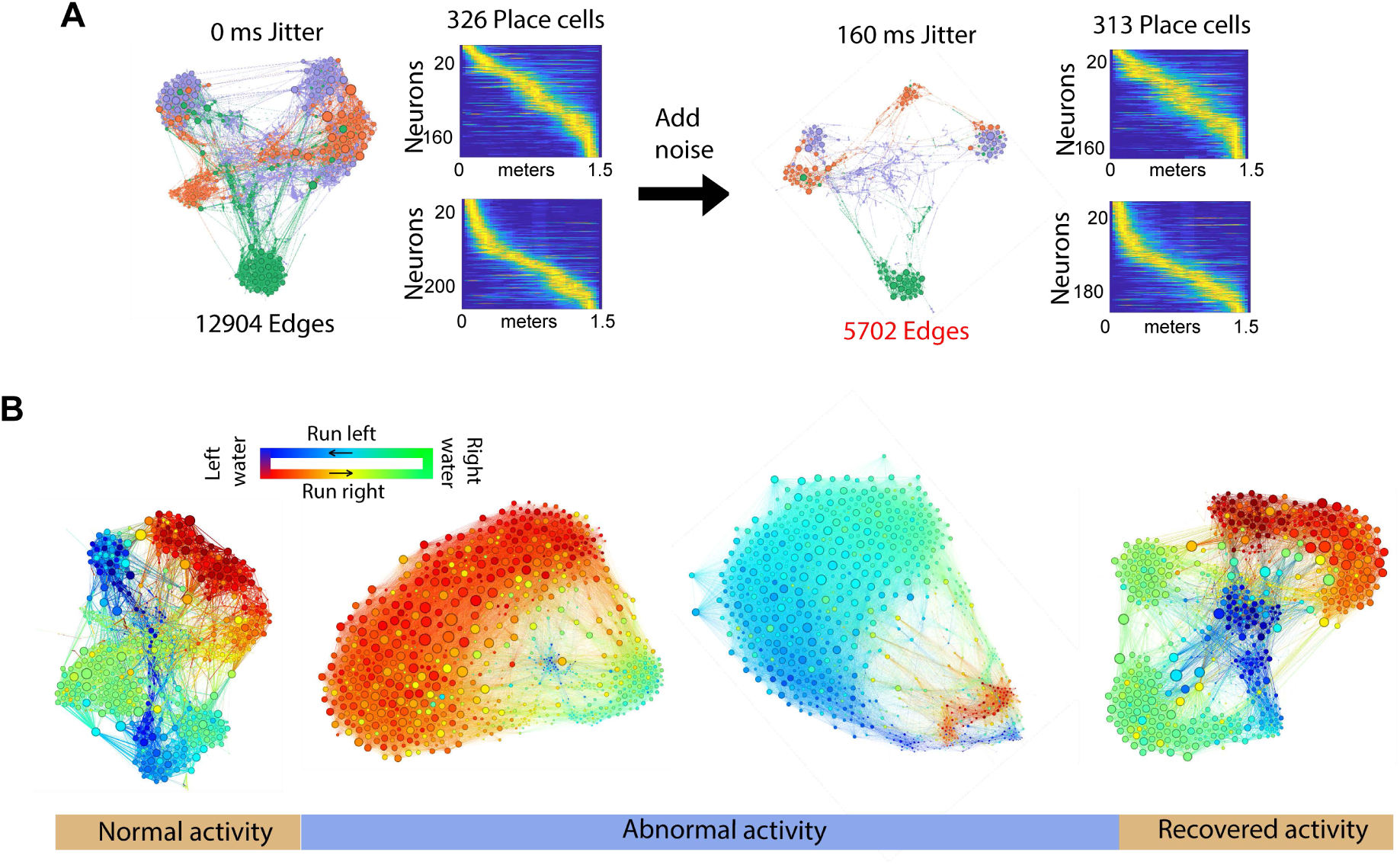
Graph topology is sensitive to noise and can capture large group of direction specific neurons after a lesion in CA1. **(A)** Introducing a small jitter noise to every deconvoluted neural spike leads to a drastic decrease (55 %) in the number of edges in a graph. However, the same amount of noise does not decrease the number of place cells detected. **(B)** Network graph during the linear track before lesions to CA1 (left), after lesion and during abnormal activity (middle), and after recovery (right). Note the large change in directional preference of the large cluster during abnormal activity.

**Figure S15.**
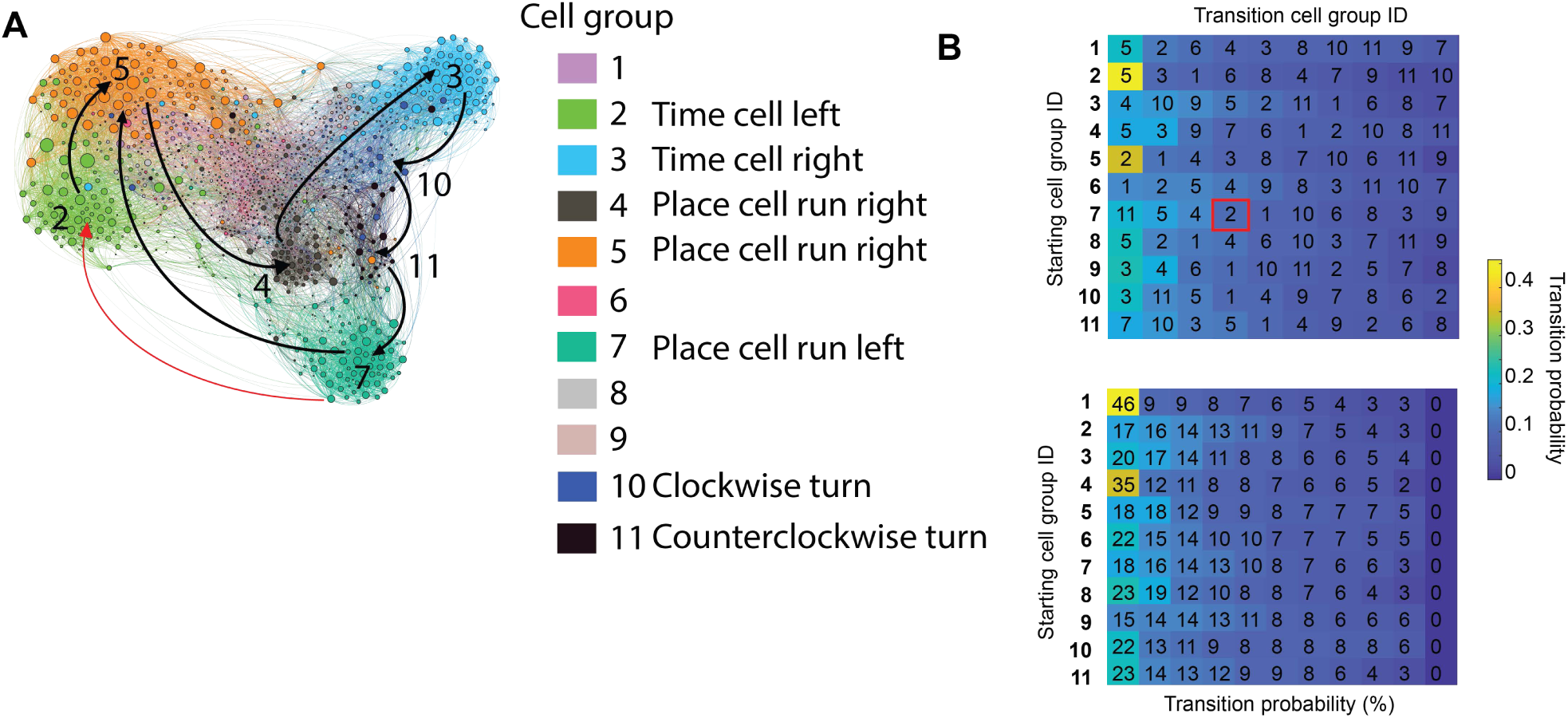
Node connectivity in a graph provides some insight into alternative sequential behaviors. **(A)** Network graph of neuronal activity of one mouse in one session. Nodes are colored by the cell group they belong to, size of the node is proportional to its degree. Arrows indicate a sequence of behaviors identified using the transition probability shown on the right panel. **(B)** The color in each box is represents the transition probability of one cell group into another (shown with numbers), the probabilities are sorted so the highest is on the left. The transition probability was calculated by counting how many edges a node makes with nodes in another cell group. Connection within the cell group are not counted. The numbers were then divided by all the edges made by that cell group, excluding self-connections. The bottom panel shows the actual probabilities shown as a percent. The graph in panel A was obtained in the following manner. Start at cell group 2, second row on right panel, the next most likely transition is to cell group 5, draw an arrow. From cell group 5, then most likely transition (ignoring loop propagation, and non-place/time cell group 1) is to cell group 4. From cell group the next transition is 3, then cell group 10 to cell group 11 and to cell group 7. From cell group 7 the next most likely transition is cell group 5, which is classified as place cell run right. This is an unlikely transition since before the animal runs right it must stop and drink. So, the only transition missing to complete the cycle is from cell group 7 to 2, which is the fourth transition (shown in red arrow and rectangle). There are many explanations for the lack of a complete cycle, including the fact that time and place cells in the same direction show little correlation due to their asymmetric population fields (Figure S13D). A different pathway could be possible if more than 11 cell assemblies would be identified using a faster Markov time. Interestingly, the transition from time cells on the right (cell group 2, light blue) to place cells in the left direction (cell group 7, green) has two intermediate cell groups (11 and 10). These two cell groups actually correspond to when the animal turns clockwise or counterclockwise (**Movie 9**).

**Figure S16.**
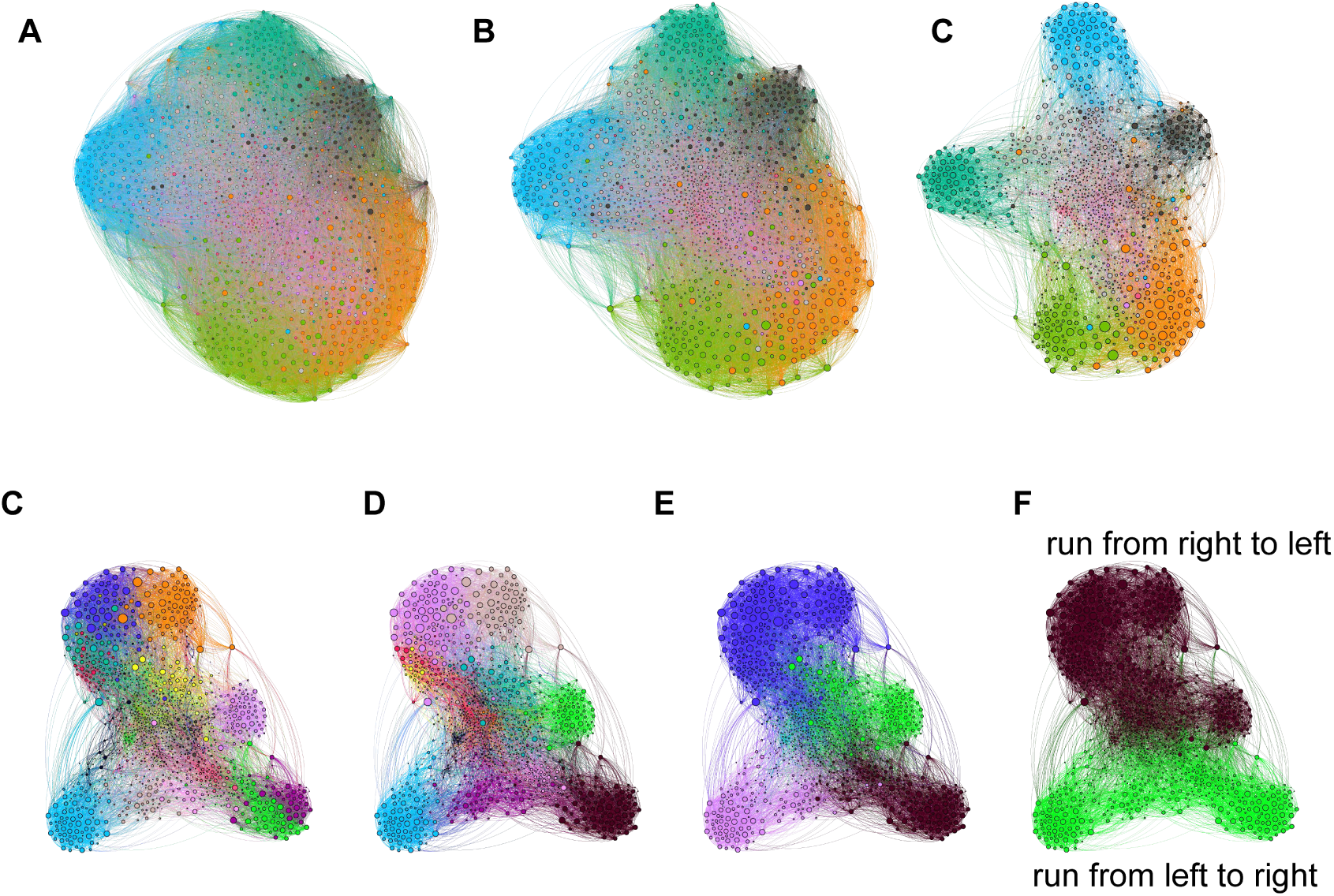
Selecting different correlation threshold or Markov diffusion time does not change the overall results presented but provide finer detail into mice behavior. **(A)** In our analysis we only look at neurons whose correlation is above 0.1 with a p value below 0.05 (bootstrap confidence calculated by MATLAB). Selecting different thresholds leads to higher or lower number of edges (left, threshold 0.0 leads to ∼ 75000 edges; middle, threshold of 0.05 leads to 35000 edges; right, a threshold of 0.1 leads to 13700 edges). The number of nodes was unchanged in the analysis. We selected a threshold of 0.1 because it provided a manageable amount of data and required a reasonable amount of time to process. However, the number of cell groups was only slightly changed (10 at correlations above 0, 11 at correlation above 0.05, and 14 at correlations above 0.1). We limited our analysis to 11 mainly because we can not assign any behavioral properties to modules beyond 6 or 7 which we know correspond to place cells. A second more important parameter is the Markov diffusion time. This parameter is proportional to how many random walks would it require for a walker to diffuse out of a cell group. **(C)** Short timescales split the graph into several groups which may correspond to finer variations of the mouse behavior in the track (time set to 0.3). Notice the time cell group being split into two. At timescales of 0.7 **(D)** and 1.0 **(E)** we start to observe a number of cell groups corresponding to direction specific place and time cells. **(F)** We noticed that longer Markov times split the graph in two groups comprising each direction in the track (time set at 5).

## Supplementary Movies

**Movie 1.** (Top row) Unprocessed 15 second recording of the left hemisphere in one animal and right hemisphere in another while running in the linear track. (Bottom row) Temporally down-sampled (4-fold) and spatially smoothed 15 second video of CA1 activity while two different mice run in the linear track. Left hemisphere is shown on the left panel and right hemisphere on the right.

**Movie 2.** Simultaneous bilateral recording of a mouse while running in the linear track. Image is background subtracted. Left hemisphere shown on the left and right hemisphere on the right. Neuronal activity extracted from this animal is shown on figure 1d.

**Movie 3.** (Left) Correlation image of CA1 activity across two months not motion corrected, each frame represents a 30-minute session (home cage exploration and linear track), 45 sessions in total. (right) The same data but motion corrected. The sudden shift between session 14 and 15 is due to the 10-day period of no task.

**Movie 4.** Motion corrected correlation image of CA1 activity in one mouse recorded for 8 months (76 sessions, data from home cage exploring and linear track). Sessions between 33 and 42 not included because abnormal activity due to damage prevent accurate registration using individual sessions. The data presented in the manuscript is motion corrected and analyzed in a different manner (see methods).

**Movie 5.** Direction specific burst activity in CA1 in three mice after damage. Note the continuously fluorescent cells, these cells will eventually become inactive. The top right animal was not included in the analysis because the lesion induced significant changes in the FOV which limited our confidence in registering cells between days. All three mice have a leftward direction burst in the video, but the direction could change between days (see figure S13).

**Movie 6.** The same data used in movie 5 but a closer view showing how some neurons persist in the field of view while other stop being active after the CA1 lesion.

**Movie 7.** (Top left) Changes in the topology of a network graph of CA1 activity in a mouse exploring its home cage across days. Note the small cluster of nodes leaving and joining the larger cluster. (Top right) Changes in the topology of a network graph of CA1 activity while the mouse becomes familiar with the linear track. (Bottom) The same data but nodes are colored by cell groups determined on the last day once the mouse is familiar with the environment. The video proceeds in reverse order, trained periods are shown in the first frames and learning are the last. Note that modules retain a large portion of neurons across days but eventually fall apart during learning. Also, the diffusion of colors indicates that some neurons can change their role in the network.

**Movie 8.** The number of edges between clusters can be used to extract the sequence of events in the behavior (see figure S14). In this video, demonstrate that a sequential behavior has a clear sequential activity in the graph. Cell groups are shown in color once at least 5 neurons in the group are active, only on cell group per frame shown.

**Movie 9.** In some cases, we observe that sequences in a graph can bifurcate, activating different groups. In this case we have at the right side of the maze 4 cell groups, cell group 3 shown in light blue (time cells), cell group 10 in dark blue, 11 in black, and 7 in green (place cells). Using the graph connectivity, we observe that the pathway 3-10-7 and 3-11-7 have very similar transition probabilities. Here we show 20 frame segments of behavior centered at when either cell group 10 (left video) or cell group 11 (right) are active. We observe that each of these sequences correspond to turning counterclockwise or clockwise.

